# Water will find its way: transport through narrow tunnels in hydrolases

**DOI:** 10.1101/2023.05.24.542065

**Authors:** Carlos Sequeiros-Borja, Aravind Selvaram Thirunavukarasu, Cedrix J. Dongmo Foumthuim, Jan Brezovsky

## Abstract

An aqueous environment is vital for life as we know it, and water is essential for nearly all biochemical processes at a molecular level. Proteins utilize water molecules in various ways. Consequently, proteins must transport water molecules across their internal network of tunnels to reach the desired action sites, either within them or functioning as molecular pipes to control cellular osmotic pressure. Despite water playing a crucial role in enzymatic activity and stability, its transport has been largely overlooked, with studies primarily focusing on water transport across membrane proteins. The transport of molecules through a protein’s tunnel network is challenging to study experimentally, making molecular dynamics simulations the most popular approach for investigating such events. In this study, we focused on the transport of water molecules across three different α/β-hydrolases: haloalkane dehalogenase, epoxide hydrolase, and lipase. Using a 5 μs adaptive simulation per system, we observed that only a few tunnels were responsible for the majority of water transport in dehalogenase, in contrast to a higher diversity of tunnels in other enzymes. Interestingly, water molecules could traverse narrow tunnels with sub-angstrom bottlenecks, which is surprising given the commonly accepted water molecule radius of 1.4 Å. Our analysis of the transport events in such narrow tunnels revealed a markedly increased number of hydrogen bonds formed between the water molecules and the protein, likely compensating for the steric penalty of the process. Overall, these commonly disregarded narrow tunnels accounted for ∼20% of the total water transport observed, emphasizing the need to surpass the standard geometrical limits on the functional tunnels to properly account for relevant transport processes. Finally, we demonstrated how the obtained insights could be applied to explain the differences in a mutant of the human soluble epoxide hydrolase associated with a higher incidence of ischemic stroke.

## Introduction

An aqueous environment is essential for life as we know it, and water is a prerequisite for nearly all biochemical processes at a molecular level. Water can influence the folding of proteins,^1–3^ their dynamics,^4^ and actively participate in chemical transformations.^5^ Proteins can utilize water molecules in various ways: as proton donors or receivers,^6^ molecular stabilizers,^7^ molecular lubricants enhancing protein dynamics,^8^ and as significant contributors behind the enthalpy– entropy compensation in protein–ligand binding.^9^ In this regard, proteins must transport water molecules across their internal network of tunnels to reach the desired action sites, either within them or as molecular pipes to control cellular osmotic pressure.

The transport of molecules through a protein’s tunnel network has proven challenging to study experimentally, with a few exceptions.^10^ Therefore, the most suitable approach to studying such events is molecular dynamics (MD) simulations, a methodology with its own limitations. Nevertheless, the principal advantage of MD simulations is that it provides a detailed atomic picture of the transport process itself, the tunnel pathways employed by the molecules, and even approximated transport rates.^11–13^ Even though water plays a crucial role in enzymatic activity and stability, its transport has been somewhat neglected, with studies mainly focused on water transport across membrane proteins like aquaporins.^14–17^

Water transport at a molecular level has been extensively studied in the nanomaterials field, where researchers have tested the peculiarities and limits of water transport through various molecular barriers.^18–21^ On the other hand, in biological sciences, the focus of water transport in proteins is primarily on the movement across membranes facilitated by specialized trans-membrane proteins, such as aquaporins. Within these proteins, the so-called single-file transport of water can be regarded as the spatial limit required for transport.^13,22,23^ Here, water molecules are arranged in a single lane and move through tunnels at least one water molecule wide, with a commonly accepted radius of 1.4 Å.^24–26^

Recent research has revealed that the methods for assessing a protein’s tunnel network and the migration of small molecules do not consistently produce equivalent results. Some water molecules could not be traced to a defined tunnel, indicating either the method’s limitations in detecting tunnels or the possibility that water molecules could traverse the protein through inconspicuous empty spaces.^27^ In this paper, to reconcile these observations, we explore the hypothesis that water transport in proteins can occur through considerably narrow tunnels with a bottleneck radius below 1.4 Å, and that this contribution is not negligible. As model systems, we focus on the tunnel network across three members of the α/β-hydrolase superfamily, emphasizing the tunnels used for water transport. Following this, we characterize these narrow tunnels and their interactions with water molecules. We demonstrate that water molecules can overcome van der Waals repulsion by an increase in the number of H-bonds. Next, to showcase the utility of the insights obtained, we analyze two variants of human epoxide hydrolase, a wild type and a mutant, resulting from a single nucleotide polymorphism (E470G), associated with an increased risk of ischemic stroke.^28^ We show that these narrow tunnels can identify structural differences in protein variants that are not easily discernible with traditional analyses. Finally, we suggest revisiting the current considerations on the parameters of functional tunnels.

## Materials and Methods

### Preparation of structures

The crystallographic structures for the systems were obtained from the Protein Data Bank^29^ (PDB): haloalkane dehalogenase (Hal) from *Rhodococcus rhodochrous*^30^ (PDB: 4E46), epoxide hydrolase I (Epx) from *Solanum tuberosum*^31^ (PDB: 2CJP), open^32^ and closed^33^ states of lipase (Lip) from *Diutina rugosa* (previously *Candida rugosa*) (PDB: 1CRL and 1TRH, respectively), and human epoxide hydrolase^34^ (hEpx) (PDB: 4X6X). In the case of Epx, two chains corresponding to two distinct enzyme structures were present in the asymmetric unit of the respective PDB record. To provide more conformational diversity, both structures underwent follow-up modeling after synchronizing their composition by removing residue K2 present only in chain A. To obtain the E470G variant of hEpx, residue E470 was renamed to glycine, and the side-chain atoms were removed from hEpx. All structures were processed by eliminating all non-protein molecules except the crystallographic waters.

The initial protonation states were determined with the H++ 3.0 webserver^35^ at pH 8.5^36^ for Hal, 6.8^37^ for Epx, 8.0^38^ for Lip, and 7.4^39^ for hEpx and E470G. For the Lip system, residue E208 was differentially protonated for the open and closed states; therefore, to build a uniform topology, this residue was manually de-protonated in both conformations. For all proteins, water molecules were initially placed around the solute using a tandem approach based on the 3D reference interaction site model theory^40^ (3D-RISM) and the Placevent^41^ algorithm. All predicted 3D-RISM waters were subsequently combined with crystallographic waters, keeping only the water molecules at least 2 Å away from the protein using EDIAscorer,^42^ a method to compute electron density for individual atoms in a crystal structure. The resulting system was then processed with the tleap module of the Amber18^43^ package. Each protein was positioned at the center of a periodic truncated octahedral box, with a distance of 10 Å away from the edges, and solvated with the 3-charge, 4-point rigid OPC water model.^44^ Na^+^ and Cl^-^ ions were initially added to neutralize the system’s charge and subsequently to reach a salt concentration of 0.1 M. The MD simulations were performed with the pmemd.cuda module of Amber18,^45,46^ employing the ff14SB^47^ force field. Finally, the hydrogen mass repartitioning method^48^ was applied to the topology to enable a simulation timestep of 4 fs.

### MD protocol

The systems underwent five minimization cycles with a stepwise release of positional restraints on the protein atoms. The minimization cycles comprised 100 steps of the steepest descent minimization algorithm, followed by 400 steps of the conjugated gradient. In the initial minimization cycle, harmonic positional restraints were imposed on all heavy atoms of the protein with a force constant of 500 kcal·mol^-1^·Å^-2^. Subsequent cycles applied restrictions only to the backbone atoms with force constants of 500, 125, 25, and 0.0001 kcal·mol^-1^·Å^-2^ sequentially. Following the minimization workflow, a brief heating round of NVT MD simulation was applied while keeping the protein’s heavy atoms restrained with a force constant of 5 kcal·mol^-1^·Å^-2^. The heating cycle involved 20 ps of MD simulation from 0 to 200 K using the Langevin thermostat with a collision frequency of 2 ps^-1^, a coupling constant of 1 ps, and a simulation timestep of 4 fs. The long-range electrostatic interactions were computed using the particle mesh Ewald summation^49,50^ and all bonds involving hydrogen atoms were constrained using the SHAKE^51^ algorithm.

Subsequently, four equilibration cycles were conducted. First, in the NVT simulation, the temperature was raised to the target value of 310 K in 100 ps, employing the same parameters as previously described. The temperature was maintained constant for 900 ps, and during this process, harmonic positional restraints were applied to the protein’s heavy atoms with a force constant of 5 kcal·mol^-1^·Å^-2^. Next, 1 ns of NPT MD simulation was run, with pressure controlled using the weak-coupling Berendsen barostat with a coupling constant of 1 ps, and positional restraints were applied only to the backbone atoms using the same force constant as in the previous stage, followed by 1 ns of NPT simulation without positional restraints. Finaly, another 200 ns of NPT simulation with the same settings as before was performed except for using Monte-Carlo barostat to enable equilibration of internal water molecules. The final snapshots from simulations of all variants of each system were utilized as their initial seeding structures for the adaptive MD simulations.

The production stage utilized the High-Throughput Molecular Dynamics^52^ package (HTMD v1.13.10) with Amber18 as the MD simulation engine. For HTMD settings, five parallel MD simulation runs per epoch were configured with 10 epochs in total. The HTMD package facilitates the construction of Markov state models (MSM) after each epoch, prioritizing less-sampled conformational states in subsequent rounds to enhance system sampling.^52–54^ To build the MSM, the dihedral angles of selected residues were employed, chosen as follows. Initially, the geometric center of the active site residues for each system was calculated. Residues N41, D106, W107, E130, and H272 for Hal; D105, Y154, Y234, D265, and H300 for Epx; G123, G124, S209, A210, E341, and H449 for Lip; and D335, Y383, Y466, and H524 for hEpx and E470G were used as centers of selection spheres (numbering according to their PDB id). The dihedral angles of all residues within a sphere of 12 Å radius for Hal, Epx, hEpx, and E470G, and 18 Å radius for Lip were used as metrics for MSM building, with a TICA^55^ dimension of 3 and a lag time of 2. The equilibration phase in HTMD comprised two short NVT and NPT MD simulations of 250 ps each. During these rounds, the systems were heated from 0 to 310 K with a Langevin thermostat and harmonic positional restraints applied to the backbone atoms with a force constant of 5 kcal·mol^-1^·Å^-2^. Finally, a 100 ns unrestrained NVT production simulation was conducted, employing the Berendsen barostat and a saving frequency of 10 ps, as only the NVT production simulations were supported by HTMD package.

### Tunnel network and water transport analyses

The trajectories underwent analysis using the cpptraj^56^ module of Amber18, CAVER^57^ v3.02, AQUA-DUCT^58^ v1.0.11, TransportTools^59^ v0.9.0, and Python3 *in-house* scripts. For the tunnel network calculations with CAVER, the divide-and-conquer approach^60^ was applied, setting the probe radius to 0.7 Å and the shell radius to 3.0 Å for Hal, Epx, hEpx, and E470G, and 5.0 Å for Lip. CAVER starting atoms for tunnel search were chosen as follows: D106-CG, W107-CD2, F168-N, and L246-N for Hal; W106-CD2, L181-N, Y235-CB, and D265-CA for Epx; G123-O, E126-CG, G342-CA, and T416-HA for Lip; Y466-CD1 and H524-CA for hEpx and E470G. The remaining parameters were left as default. These atoms were selected based on their center of geometry, presenting one of the lowest root mean square fluctuations (RMSF), and being inside the deepest part of the cavities described in the literature for each protein.^61–70^ To achieve consistent tunnel entrances, HDBSCAN v0.8.27^71^ was used at the tunnel endpoints for clustering, with the following parameters: *cluster_selection_epsilon* of 1.5, *allow_single_cluster* set to True, *min_samples* and *min_cluster_size* set to 5. The formed clusters were considered independent tunnels, and the tunnels identified as noise were discarded. For water tracking analyses, the same atoms described earlier were used for the object definition in AQUA-DUCT, employing a sphere of 6.0 Å for Hal and 4.0 Å for Epx, Lip, hEpx, and E470G. The remaining parameters were set to default. In TransportTools, the average-link hierarchical clustering method with a threshold of 1.0 Å was applied, utilizing the exact matching analysis for the assignment of transport events. All other settings were maintained as default.

For each water transport event assigned by TransportTools to a defined tunnel for at least 70% of its duration, a post-processing step was performed to obtain more detailed information about the nature of the transport. Here, a tunnel is regarded as a collection of *tunnel spheres* along a central line, and in each frame of a transport event, the water molecule involved is assigned to the closest *tunnel sphere* (**Figure 1**). Following this concept, a transport event comprises a collection of *event spheres* to which a migrating water molecule was assigned, with a restriction of a single *event sphere* per frame. Our subsequent analyses focused on frames where the water molecule was assigned to the smallest sphere along the event, referred to as the *minimal event sphere* (**Figure 1**). For these selected frames, the radius of the *minimal event sphere* and the hydrogen bonds (H-bonds) between the water molecule and the protein were considered. Energetically significant H-bonds with a distance between donor and acceptor heavy atoms up to 3.5 Å were identified using the cpptraj program.^72^

**Figure 1.**
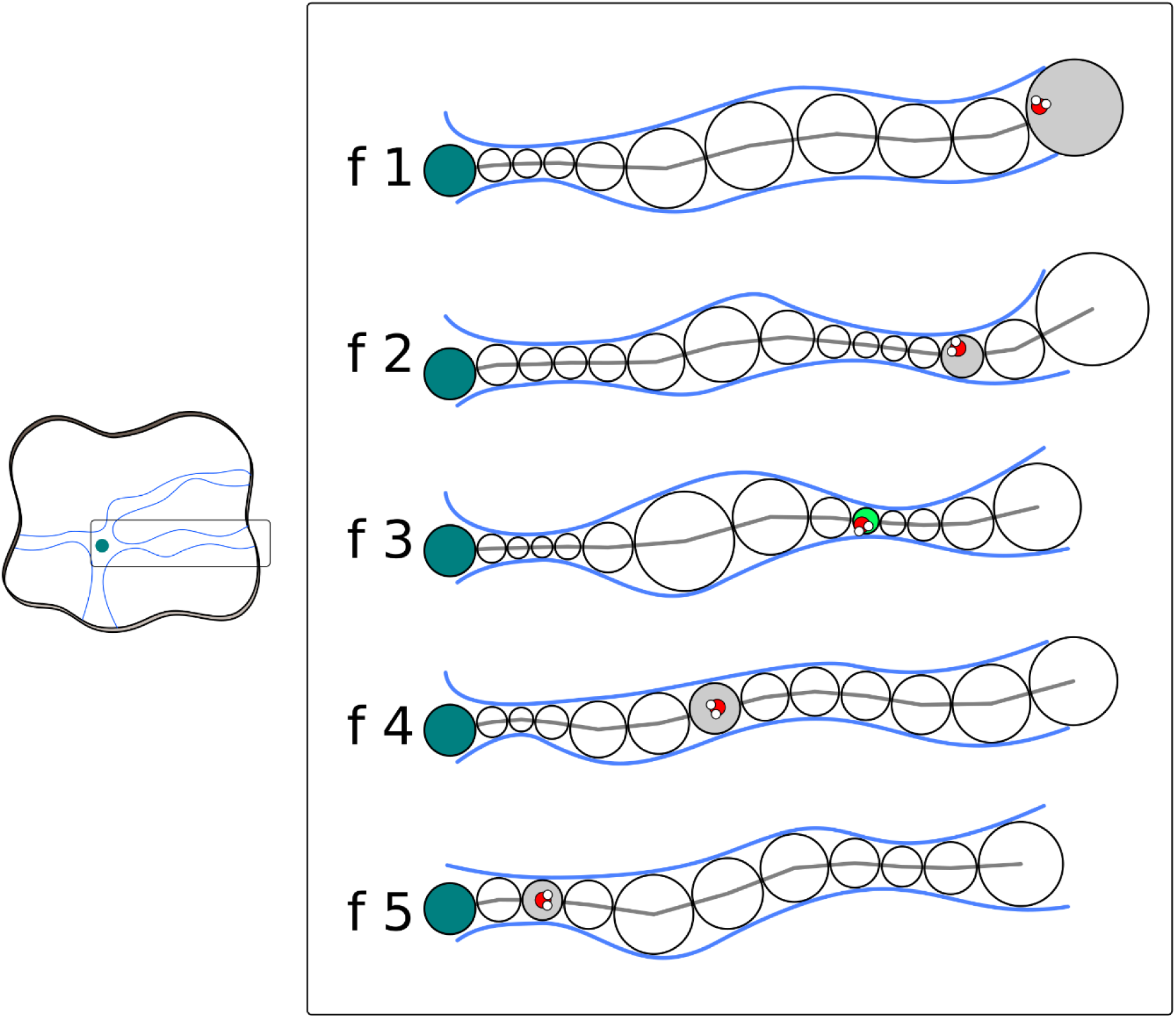
Water transport event and *minimal event sphere* definition. A protein can contain several tunnels and transport events (left), and in one tunnel, a transport event is composed of several frames (enclosed right). In each frame of a transport event, a tunnel is present as a set of *tunnel spheres* (empty circles) along a centerline (gray line) that joins the active site (turquoise circles) with the exterior. In each frame, the migrating water molecules are assigned to a single *event sphere* of the tunnel (gray circles), and among all of them, the smallest corresponds to the *minimal event sphere* (green circle).

## Results and Discussion

### Tunnel networks identified in Hal, Epx, and Lip from adaptive MD simulations

The MD simulations for all the proteins exhibited stable behavior during the 5 μs of simulation, with root mean square deviation (RMSD) averages of 1, 2, and 2.5 Å for Hal, Epx, and Lip, respectively (**Figures S1–S3**). Although these differences in RMSD might raise some concerns about the stability of the systems, the RMSF analyses showed that this is not the case. The size variations in proteins are different, and in Hal, the protein is densely packed, with fluctuations mainly occurring at the N- and C-terminal regions (**Figure S4**). Similarly, in Epx and Lip, some regions show higher flexibility than the rest of the system (**Figures S5 and S6**), corresponding to areas with large b-factors in the crystals, indicating high flexibility.

TransportTools results for Hal showed the presence of all tunnels described in the literature^61^, namely *p1*, *p2a*, *p2b*, *p2c*, and *p3*^62^ (**Figure S7a,b and Table S1**), with all of them used for water transport to some extent. The results also revealed new tunnels for water transport: a top tunnel near *p1* (called the *up* tunnel here), located closer to helix α5’ (**Figure S7c,d** – salmon color), and a side tunnel perpendicular to the *p1* tunnel, opposite to the *p2* tunnel (called the *side* tunnel here), between the α/β and cap domains (**Figure S7c,d -** light green color). For Epx, the three main tunnels described by Mitusińska *et al.* are present,^63^ but tunnel *TM1* is formed by two TransportTools superclusters (**Figure S8a,b and Table S1**). The results also show numerous new tunnels that transport water, albeit in smaller quantities, which may correspond to the outliers observed by Mitusińska *et al.* (**Figures S8c,d and S9**). Lip results showed predominant water transport in the *main* tunnel described in previous studies^64–67^ (**Figure S10a,b and Table S1**). Additionally, several important tunnels for water transport were identified, most in the same region of the main tunnel (**Figure S10c,d**) and fewer in different regions (**Figure S11**). The results also revealed the presence of the hypothesized *ester* exit tunnel;^68^ however, only two water molecules were transported via this tunnel (**Figure S10a,b and Table S1**). Tunnel network analysis for the three systems showed the presence of all tunnels described in the literature, with several new tunnels found, mainly for Epx and Lip.

### Water transport within the tunnel networks of Hal, Epx, and Lip

Overall, we collected 2222, 17786, and 11086 water transport events that could be unambiguously assigned to tunnels in Hal, Epx, and Lip, respectively (**Table S2**), constituting 23–84% of all transport events analyzed. A limited number of events (<3%) could not be assigned even to the overall superclusters generated by TransportTools across all 50 simulations of each hydrolase. However, the main reason for misalignment between water events and tunnels originated from attempts to match these events exactly to particular tunnels found by CAVER in the trajectory part where the transport event occurred, which is in line with recent observations for eight soluble epoxide hydrolases.^27^ In general, mismatches stem from the approximation of asymmetric internal voids by spherical tunnels produced by CAVER,^73^ promoted by maintaining only the tunnel with the best throughput for the tractability of tunnel clustering in each frame,^74^ restricting the description of the entire tunnel geometry.

For the assigned events, the distribution of *minimal event sphere* radii for Hal exhibited two peaks, with the major peak at 1.6–1.7 Å and a minor peak at 1.1–1.2 Å (**Figure 2**). A similar trend was observed for Epx, with a minor peak at 1.2–1.3 Å and a major one at 2.2–2.3 Å (**Figure 2**). However, this trend changed in Lip, where the curve showed a clear single peak at 1.5–1.6 Å (**Figure 2**). An examination of these distributions by the tunnels involved revealed that the bimodal nature of Epx transport was primarily due to two tunnels (**Figure S12**). A somewhat similar phenomenon was observed for Hal, in which two tunnels formed a smaller peak around 1.0–1.2 Å (**Figure S12**). This bimodal behavior was not the case in Lip, because all tunnels featured a peak at 1.5–1.6 Å (**Figure S12**). Cumulative distribution results displayed a sigmoidal behavior for all systems, although Epx deviated slightly due to its stronger bimodal nature (**Figure 2**). Given 1.4 Å as the widely accepted radius for a water molecule,^24–26^ the number of transport events below this threshold was 22.7%, 17.7%, and 25.0% for Hal, Epx, and Lip, respectively (**Figure 2**), not a negligible quantity at all.

**Figure 2.**
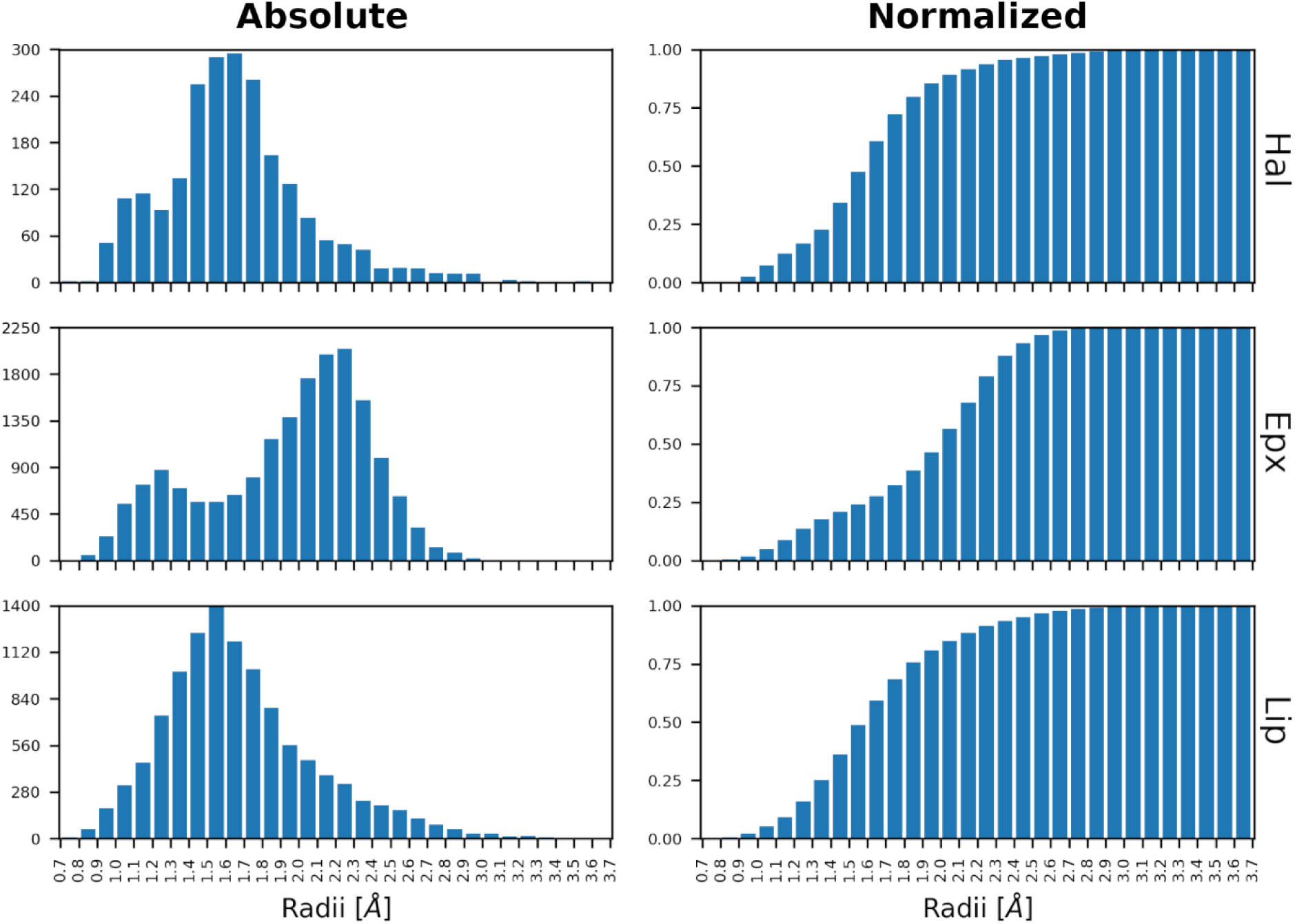
Distribution of *minimal event sphere radii* for water transport in Hal, Epx, and Lip. The absolute number of transport events and their corresponding normalized cumulative distribution.

Before we considered the implications of these observations, we had tested that such narrow *minimal event sphere* radii were not due to well-known limitations arising from the spherical approximation of transport tunnels by CAVER.^73^ Such approximation could overlook available voids in the protein structure adjacent to the tunnel sphere that could easily accommodate the migrating water molecule. By measuring the surface-surface distances between water molecules in the *minimal event sphere* and the closest protein atoms (**Figure S13)**, we verified that water molecules were indeed passing through restricted empty spaces, marked by frequent atomic overlaps for most of these events.

For proteins specialized in water transport, such as aquaporins, the accepted minimum radius of a functional state is the size of a water molecule, leading to the so-called single-file water transport.^22^ Given the results obtained for the three hydrolases studied here, it becomes evident that water can often be transported below the limits recognized for such optimized proteins, where the optimized flow of water molecules is critical to maintaining cellular function. Intrigued by these results, we focused on unraveling the characteristics and limits of water transport through enzyme tunnels with radii below 1.4 Å.

### Water transport through narrow tunnels

We examined H-bonds between the protein and water for each frame in which the water molecule was inside the *minimal event sphere*. Notably, water demonstrated a tendency to form several H-bonds with the protein at a lower *minimal event sphere* radius, a consistent pattern observed across the three systems studied here (**Figure 3**). This trend persisted when considering all radii for the analysis and was further emphasized by an apparent inverse correlation between the radius and the number of H-bonds (**Figure S14**).

**Figure 3.**
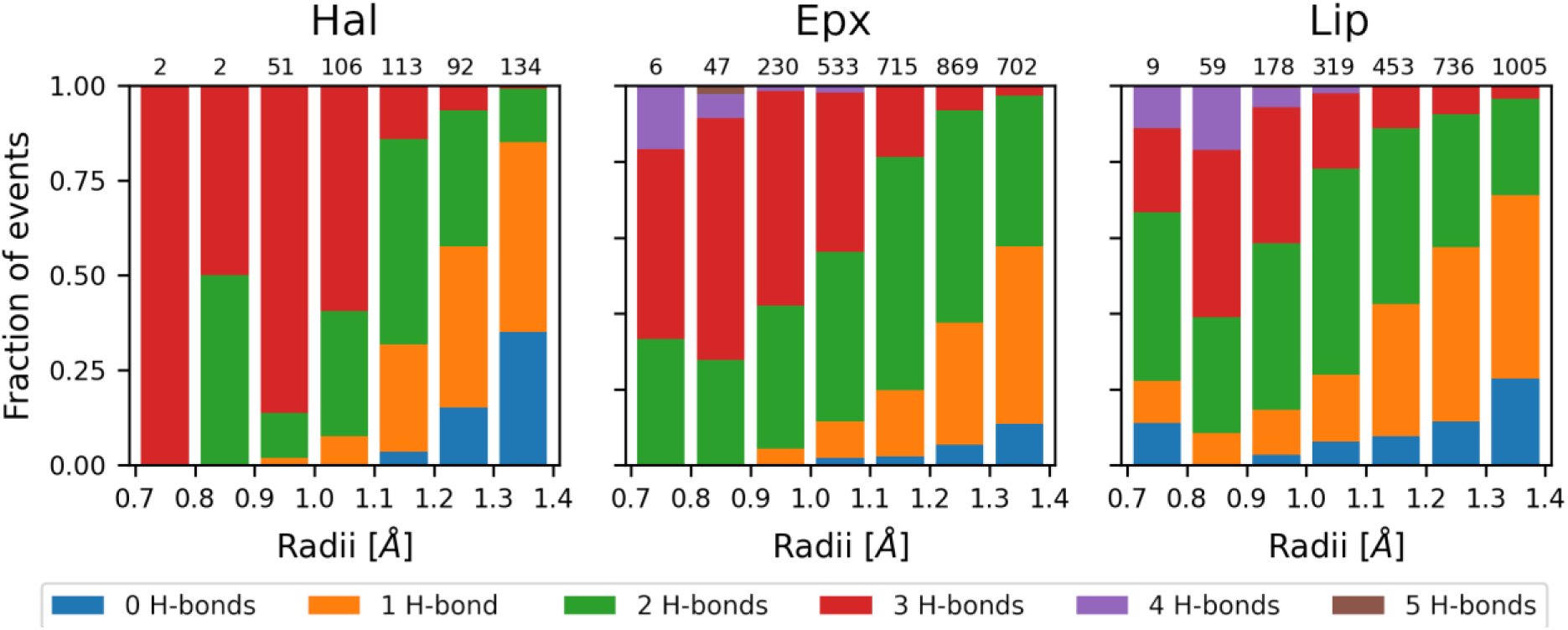
Hydrogen bond distribution of water events at narrow tunnels. H-bonds formed between water and protein at narrow tunnels with bottleneck radii below 1.4 Å. The normalized distributions for Hal, Epx, and Lip are shown, and the total number of events in each bin is displayed at the top.

Our hypothesis suggests that water molecules require multiple H-bonds to stabilize their transport through narrow tunnels, enabling them to overcome repulsive van der Waals forces between water and protein atoms (**Figure 4**). This behavior aligns with a mutational study on the trans-membrane protein 175 from *Chamaesiphon minutus*,^26^ where an increase in wetting behavior at the gate was observed, despite the tunnel radius remaining constant. The mutation involved changing hydrophobic gate residues Ile and Leu to the polar Asn. While this observation pertains to wetting rather than transport, it aligns with our findings. Our H-bond analysis contradicts the notion of single-file water transport, where the presence of H-bond donors/acceptors along the entire tunnel length is believed to increase the energy barrier for water transport.^22,23^ In the realm of nanomaterials, a pore radius of 2.85 Å in a graphdiyne membrane was considered the ultimate size limit for effective water transport.^20^ Despite being almost twice the size of a water molecule, the entirely hydrophobic nature of these artificial structures prevents water from physically passing through the open pores.^18^

**Figure 4.**
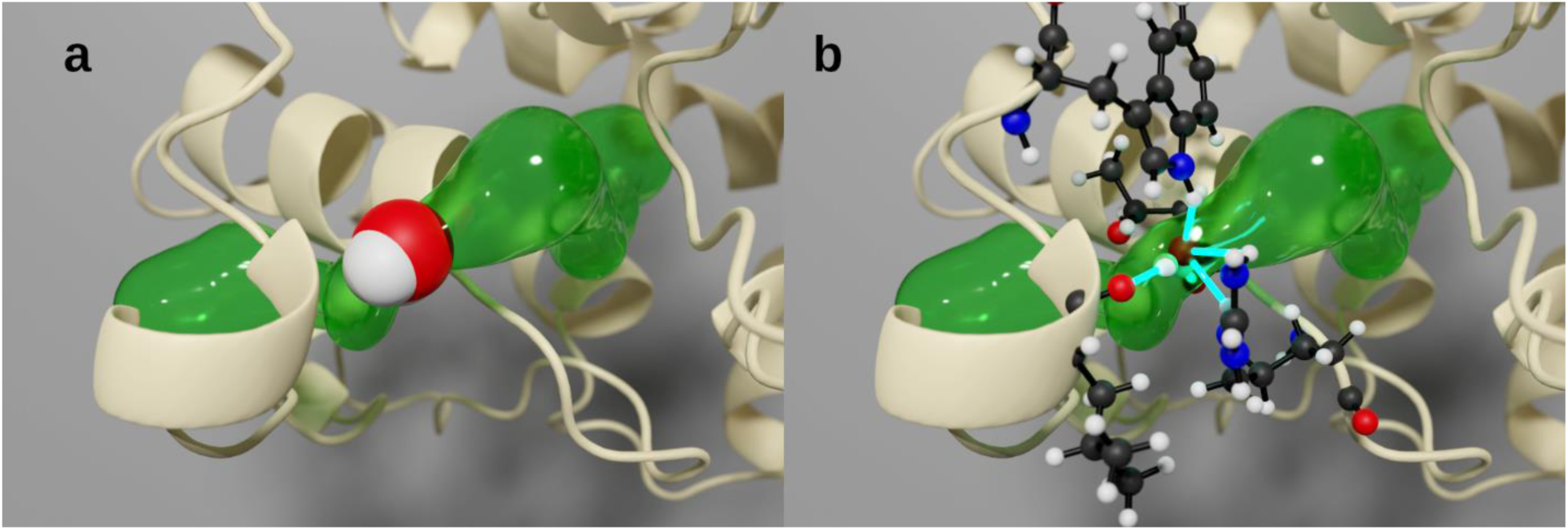
A water molecule forming hydrogen bonds while traversing a narrow tunnel. (a) A water molecule (van der Waals volume) inside Hal (beige cartoon), traversing a narrow tunnel (green surface). The volume of the water molecule exceeds the available space in the tunnel. (b) An alternative representation of the same frame, with the water molecule (depicted as ball and sticks) inside Hal (beige cartoon), forming multiple H-bonds (cyan sticks) with protein sidechain and backbone atoms (ball and sticks) during its passage through a narrow tunnel (green surface).

Furthermore, our analysis of which atoms on the residues were responsible for the H-bonding network revealed a predominance of the backbone atoms (**Figures 5 and S15**). Given that each H-bond analyzed could involve multiple atoms making contacts, the distributions visibly changed for Hal and Epx, not only in quantity but also in the position of the peaks (**Figure S15**). In the Lip case, only the quantity changed, maintaining the peak almost at the same radius (**Figure S15**). Although there is no clear trend regarding the type of atoms forming H-bonds with water during transport across all enzymes, it is reasonable to conclude that backbone atoms play a crucial role, especially at lower radii. This insight leads to the hypothesis that water transport through considerably narrow tunnels is fold-specific and may be preserved within protein families. Supporting this observation, a recent study on epoxide hydrolases demonstrated that tunnels are conserved structural features of proteins.^75^ Although mutational studies on tunnels have indicated changes in specificity^76^ and catalytic rate,^77^ the targets were primarily the major tunnels, and the observed effects were related to substrate/product intake and release.

**Figure 5.**
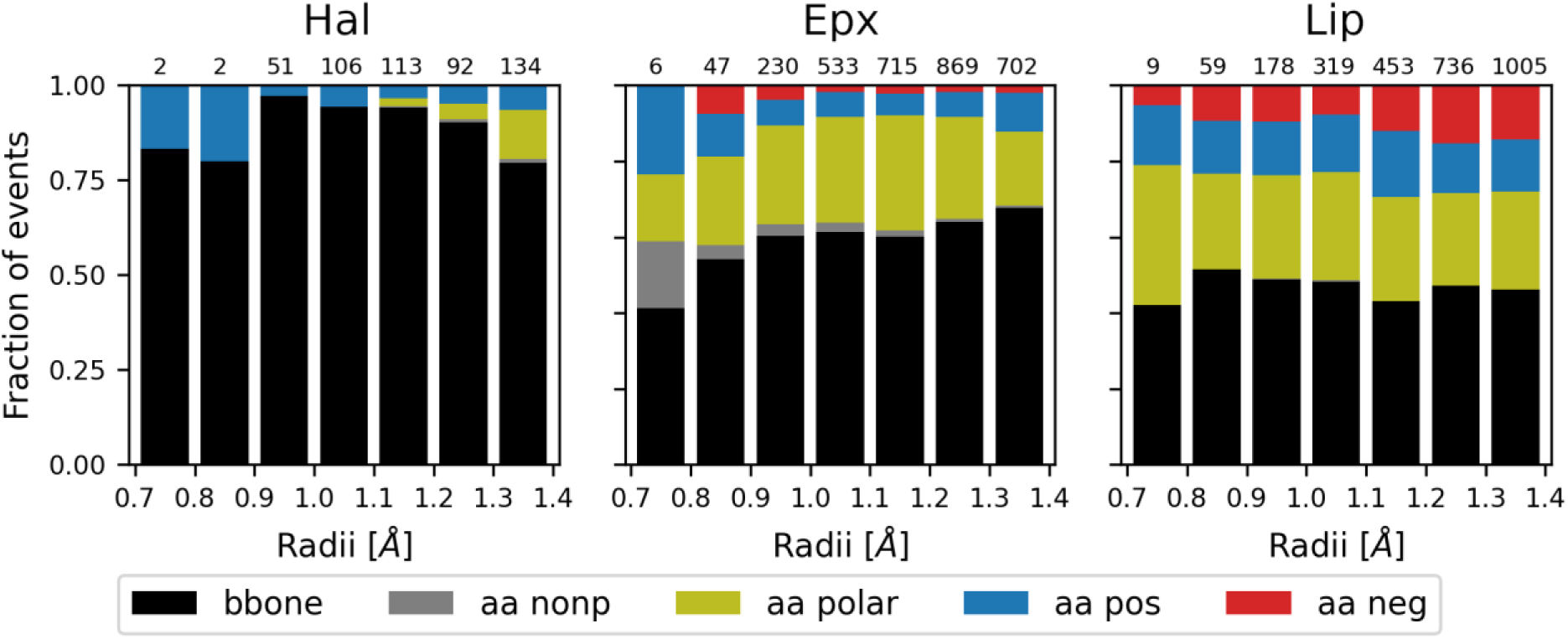
Distribution of atoms responsible for hydrogen bonding in water transport. The protein atoms involved in H-bonding with water in transport events for Hal, Epx, and Lip. Sidechain atoms were considered non-polar (Ala, Gly, Ile, Leu, Met, Phe, Pro, Trp, Val), polar (Asn, Cys, Gln, Ser, Thr, Tyr), positive (Arg, His, Lys), or negative (Asp, Glu) charged. The total number of events for each bin is indicated at the top.

### Case study: human epoxide hydrolase and risk of ischemic stroke

To illustrate the often overlooked significance of narrow tunnels, a comparative analysis was conducted between the human soluble epoxide hydrolase wild type (hEpx) and the variant E470G. hEpx utilizes water as a co-substrate for the hydrolysis of epoxides into their corresponding diols, making water a crucial component for its proper functioning.^78,79^ hEpx is involved in the hydrolysis of epoxyeicosatrienoic acids, which possess vasodilator, anti-inflammatory, analgesic, antifibrotic, and antihypertensive properties.^80^ Polymorphisms in the gene encoding hEpx have been linked to varying enzyme activity rates,^39^ thereby associated with susceptibility or protection against incident ischemic stroke.^28^ Notably, one such variant, E470G, is correlated with an increased risk of ischemic stroke in the African-American population, although the underlying molecular mechanism remains unclear.^28^

Results from 5 μs MD simulations for each variant (hEpx and E470G) revealed minimal average differences in RMSD (**Figure 6a**) across all simulations (**Figures S16 and S17**). Similarly, the average RMSF values for both variants exhibited negligible differences, except for the region spanning S407–E446 (**Figure 6b**), corresponding to a loop region on the cap domain of the protein (**Figure 6c**). This region demonstrated significantly increased flexibility in E470G simulations compared to hEpx (**Figures S18 and S19**), indicating enhanced flexibility resulting from the mutation, despite the mutation not occurring within this region (**Figure 6b,c**). The distribution of *minimal event sphere* radii for hEpx and E470G shows a bell-shaped distribution, peaking at ∼1.6 Å in both cases (**Figure S20**). On the cumulative distribution of events, the proportion of events below 1.4 Å decreased to 12.4% and 13.8% in hEpx and E470G, respectively (**Figure S20**), compared with Hal, Epx, and Lip. This reduction in transport events may be attributed to a substantial increase in total transport events between proteins (30 times higher compared to Hal and hEpx). The distribution of events by tunnels revealed that some tunnels preferred narrow radii, shifting the distribution away from the major tunnel (**Figure S21**), although not as pronounced as observed for Epx or Hal (**Figure S12**). H-bonding analyses exhibited similar patterns and trends observed in Hal, Epx, and Lip, with a predominance of H-bonds at lower *minimal event sphere* radii (**Figure S22**), and the backbone atoms being the major contributors to such contacts (**Figure S23**).

**Figure 6.**
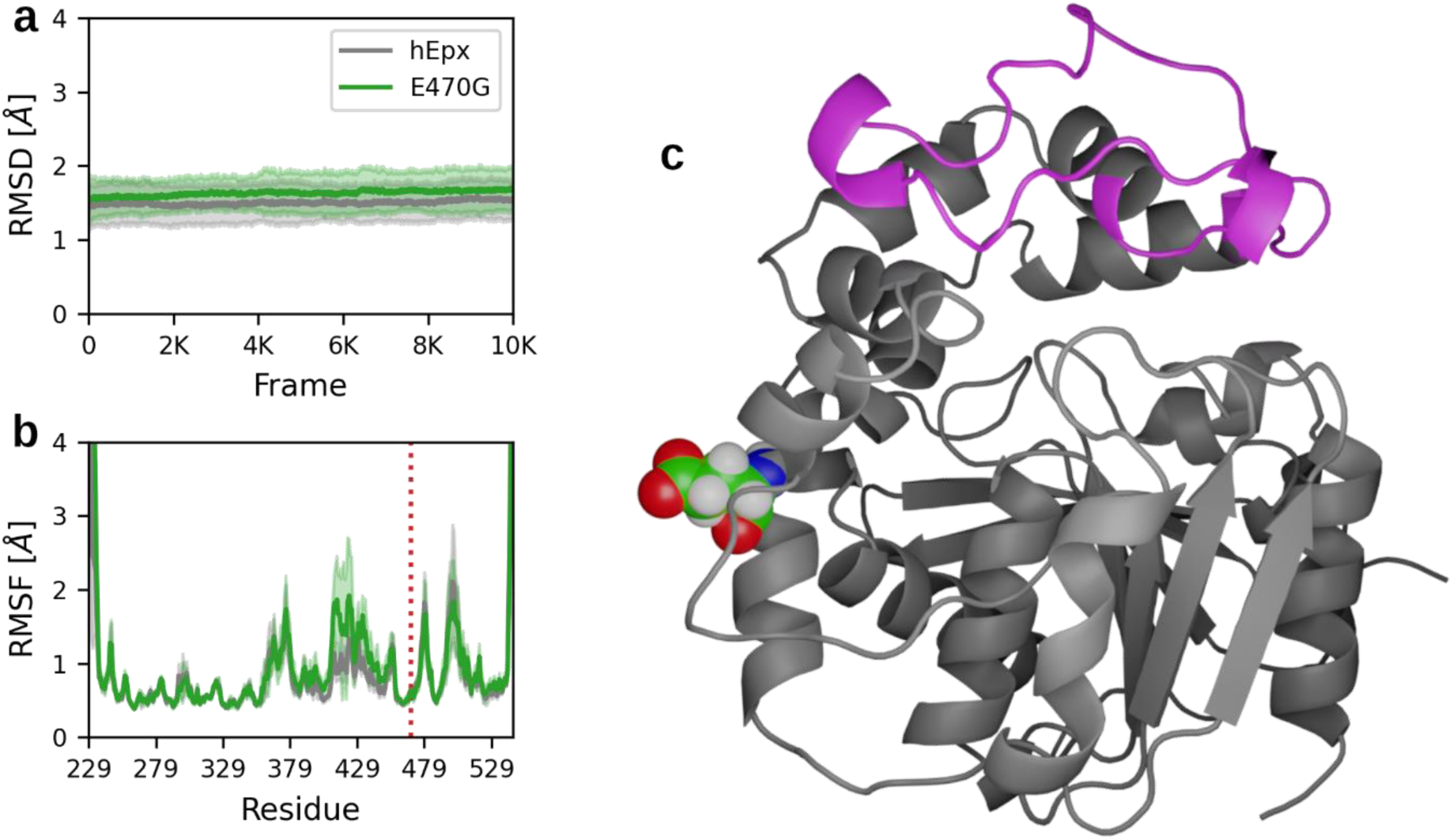
Stability and flexibility of human epoxide hydrolase wild type and E470G mutant. Average root-mean-square deviation (a) and root-mean-square fluctuation (b) are depicted as solid lines, with standard deviation represented by shaded regions for 5 μs adaptive molecular dynamics simulations. The position of the mutation is indicated by a vertical dotted red line in b. (c) Cartoon representation of hEpx with the most flexible region in magenta and the mutated residue represented as spheres.

The MD simulation results encompass all tunnels in hEpx documented in the literature,^63,69,70^ along with four newly identified ones (**Table 1 and Figures S24–S26**). When focusing solely on tunnels utilized for water transport, the *Tc/m* tunnel emerged as the major contributor in both variants (**Figure 7a and Table S3**), although its significance decreased by ∼16% in E470G compared to hEpx. Intriguingly, in a recent study of epoxide hydrolases,^70^ the *Tm1* tunnel was identified as the primary contributor to water transport in hEpx. However, this discrepancy may result from shorter simulation time and the water model employed (TIP3P versus OPC).^81,82^

**Figure 7.**
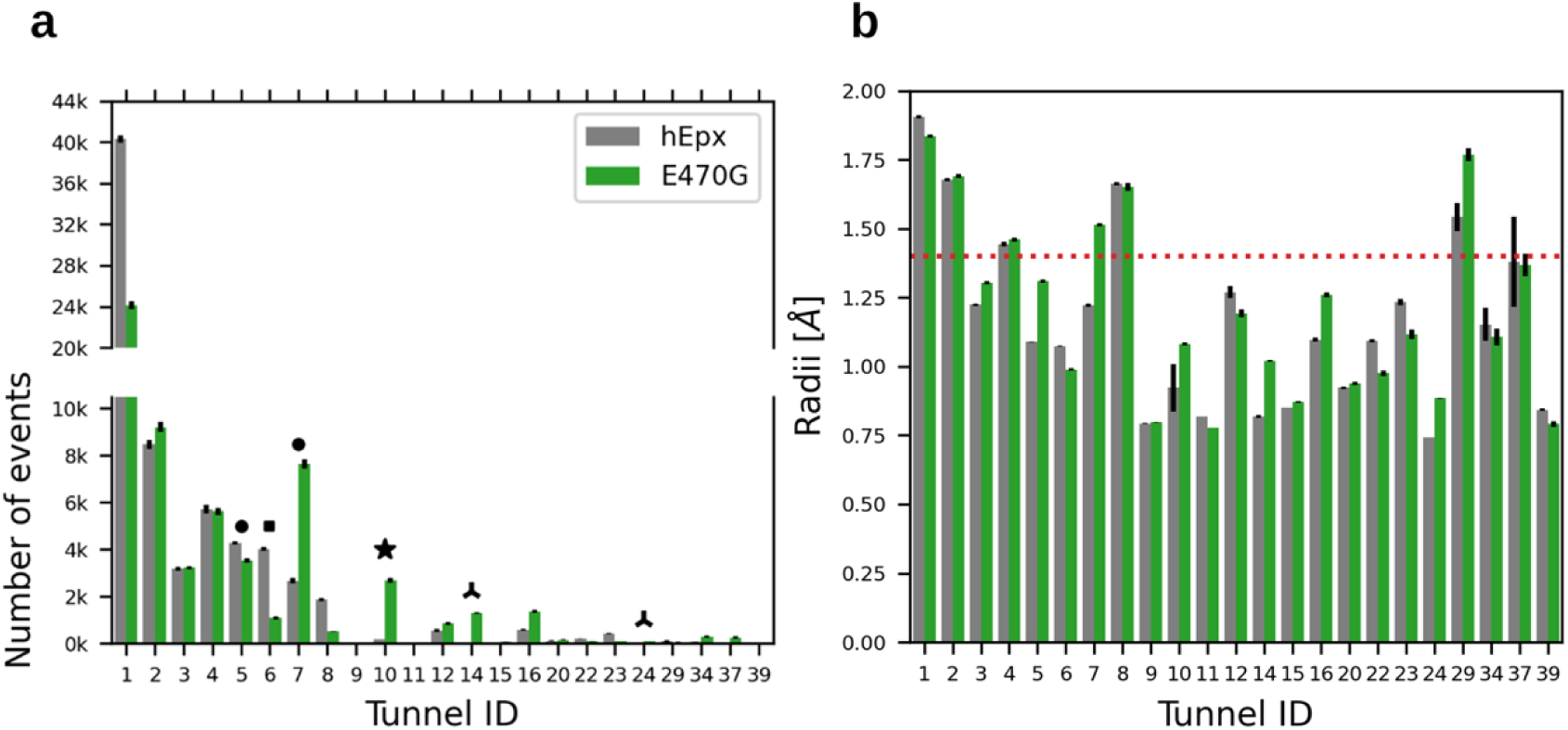
Tunnels employed by water in human epoxide hydrolase wild type and E470G mutant. (a) Total number of transport events by the tunnel for hEpx and E470G. (b) Average bottleneck radius for tunnels employed by water in hEpx and E470G, with the accepted radius for a water molecule marked by a horizontal red dotted line. Narrow tunnels with altered water transport are denoted by black shapes corresponding to the naming in **Table 1**. The reported variances were estimated using jackknife resampling from 50 contributing simulations.^83^

**Table 1.** Utilization of tunnels found in hEpx and E470G mutant.

| Tunnel | Tunnel IDs | Water transport |  |  |
| --- | --- | --- | --- | --- |
|  |  | WT | E470G | Difference |
| *Tm1* | 2, 4, 8 | 22.37% | 24.48% | -2.12% |
| *Tm2* | 16 | 0.83% | 2.45% | -1.62% |
| *Tm3* | 3, 20, 37 | 4.66% | 5.90% | -1.25% |
| *Tm5* ■ | 6 | 5.14% | 1.66% | **3.49%** |
| *Tc/m* | 1, 29, 34 | 55.86% | 39.63% | 16.22% |
| *Tg* | 12 | 0.76% | 1.40% | -0.64% |
| *Tside* ● | 5, 7 | 9.13% | 17.49% | **-8.37%** |
| *Tcap1* ▲ | 14, 24 | 0.00% | 2.27% | **-2.27%** |
| *Tcap2* | 15 | 0.01% | 0.10% | -0.08% |
| *Tcap4* | 9, 39 | 0.09% | 0.04% | 0.05% |
| *Tnew_A* ★ | 10 | 0.21% | 4.20% | **-3.98%** |
| *Tnew_B* | 11 | 0.03% | 0.02% | 0.01% |
| *Tnew_C* | 22 | 0.28% | 0.16% | 0.12% |
| *Tnew_D* | 23 | 0.62% | 0.19% | 0.43% |

An examination of the average bottleneck radius of the tunnels reveals that most have a radius below the widely accepted 1.4 Å radius for a water molecule (**Figure 7b and Table S3**). Although it is expected that the *Tm1* and *Tc/m* tunnels, with larger bottleneck radii, transport approximately 78% and 64% of total water in hEpx and E470G, respectively (**Tables 1 and S3**), the contribution of narrower tunnels should not be dismissed, particularly in E470G, where the mutation appears to have significantly altered tunnel usage (**Tables 1 and S3**). In nearly all tunnels, the volume of transported waters increased in E470G compared to hEpx (**Table S3**), likely due to reduced transport via the *Tc/m* tunnel (**Figure 7a and Table 1**). A noteworthy case resulting from the mutation is the newly identified *Tnew_A* tunnel, whose bottleneck radius increased by 0.128 Å, while its importance for water transport increased 20-fold (**Table 1 and Table S3**). The mutation also opened the *Tcap1* tunnel and increased the flow in the *Tside* tunnel in E470G (**Figure 7a and Table 1**). Similarly, a minimal reduction in the bottleneck radius of the *Tm5* tunnel by 0.085 Å in the E470G mutant led to a nearly 25% decrease in transport compared to the wild type hEpx (**Figure 7a and Table 1**).

We hypothesize that the modified water transport and tunnel usage by hEpx and E470G may contribute to the phenotypic changes observed in these variants. The reduced water flow in E470G could create a more suitable environment for catalysis, potentially leading to the increased activity observed in experimental tests.^39^ Although a comprehensive structural explanation for the observed differences in these two variants is beyond the scope of this paper, we propose that the discrepancies in water flow could offer insights into understanding their phenotypes. Moreover, the entry of a few water molecules into the active site can significantly enhance the overall efficiency of LinB86 dehalogenase,^84^ where the new inflow of water molecules through the engineered narrow tunnel notably facilitates the rate-limiting product release. It is important to clarify that differences in water transport between hEpx and E470G cannot be considered a direct cause of an increased risk of ischemic stroke. However, these differences provide more nuanced insights than a simple conformational analysis demonstrating increased flexibility in certain regions. Furthermore, variations in tunnel usage were identified in tunnels with bottleneck radii that might be easily overlooked if the focus is solely on wider tunnels. These findings also expand potential targets for protein engineering, where the opening or closing of tunnels has been successfully employed to alter relevant functional properties of enzymes.^84^

## Conclusions

In this paper, we have demonstrated that water molecules can traverse extremely narrow tunnels, even with sub-angstrom radii. Approximately 20% of the events in the evaluated proteins involved such narrow tunnels, reconciling the discrepancy between geometry-based and ligand-tracking methods for tunnel identification.^27^ We also illustrated that to overcome repulsion during transport, water molecules establish multiple H-bonds, primarily with the backbone. Furthermore, we observed that the number of H-bonds formed is inversely proportional to the *minimal event sphere* radius. Although we currently lack sufficient data to support the extrapolation of these observations beyond the α/β-hydrolase fold, our findings suggest a need to reconsider classical limits on the functional tunnels in enzymes, moving beyond the pure dimensions of their bottleneck. Lastly, based on the insights gained, we propose a structural and functional hypothesis for the observed differences in hEpx and a single-point mutant variant. These differences can be traced to minor alterations in tunnel dimensions, resulting in a major rewiring of water transport flows through the active site.

## ASSOCIATED CONTENT

### Supporting Information

The following files are available free of charge. RMSD and RMSF plots for adaptive simulations of Hal, Epx, Lip, hEpx, and E470G.Visual representation of tunnels identified in adaptive simulations of Hal, Epx, Lip, hEpx, and E470G. Distribution of *minimal event sphere* radii in Hal, Epx, Lip, hEpx, and E470G concerning tunnels, H-bonds, and interacting atoms. The closest surface-surface distances between protein atoms and water molecules present in individual bins of *minimal event sphere* radii in Hal, Epx, and Lip. Distribution of *minimal event sphere* radii in hEpx and E470G. Literature-based naming of tunnels identified in Hal, Epx, and Lip. Number of events detected and assigned to tunnels in Hal, Epx, and Lip. Water usage and average bottlenecks of tunnels in hEpx and E470G (PDF).

### Data and Software Availability

The underlying data for this study are available in the published article, the Supporting Information, and in Zenodo repositories at https://doi.org/10.5281/zenodo.7966082 (Hal), https://doi.org/10.5281/zenodo.7966059 (Epx), https://doi.org/10.5281/zenodo.7966092 (Lip), https://doi.org/10.5281/zenodo.7965661 (hEpx), and https://doi.org/10.5281/zenodo.7965881 (E470G). The data include Python3 scripts, binary files, plain text, and PDB-formatted and AMBER-formatted structural data, all compatible with various freely available SW packages. No tools with restricted access are required.

## AUTHOR INFORMATION

### Author Contributions

C.S-B. set up, performed, and analyzed simulations of hEpx and E470G, analyzed water transport in all five systems, interpreted data, and prepared the figures and manuscript draft. A.S.T. set up, performed, and analyzed simulations of Hal. C.J.D.F. set up, performed, and analyzed simulations of Lip. J.B. conceived and coordinated the project, designed the calculation, set up, performed, and analyzed simulations of Epx, and interpreted data. The manuscript was collaboratively written by all authors, who have given approval to its final version.

### Notes

The authors declare no competing financial interests.

## ACKNOWLEDGMENT

This work was supported by the National Science Centre, Poland (grant no. 2017/26/E/NZ1/00548). Computations were performed at the Poznan Supercomputing and Networking Center. C.S-B. and A.S.T. are recipients of a scholarship associated with the POWER project (grant nos. POWR.03.02.00-00-I022/16 and POWR.03.02.00-00-I006/17, respectively).

## ABBREVIATIONS

Hal: haloalkane dehalogenase from *Rhodococcus rhodochrous*
Epx: potato epoxide hydrolase 1
Lip: lipase from *Diutina rugosa* (previously *Candida rugosa*)
MD: molecular dynamics
hEpx: human epoxide hydrolase
E470G: E470G mutant of human epoxide hydrolase
MSM: Markov State Models
RMSD: root mean square deviation
RMSF: root mean square fluctuation.

## REFERENCES

(1) Levy, Y.; Onuchic, J. N. Water Mediaton in Protein Folding and Molecular Recognition. Annu. Rev. Biophys. Biomol. Struct. 2006, 35 (1), 389–415. 10.1146/annurev.biophys.35.040405.102134.

(2) Rhee, Y. M.; Sorin, E. J.; Jayachandran, G.; Lindahl, E.; Pande, V. S. Simulations of the Role of Water in the Protein-Folding Mechanism. Proc. Natl. Acad. Sci. 2004, 101 (17), 6456–6461. 10.1073/pnas.0307898101.

(3) Dongmo Foumthuim, C. J.; Giacometti, A. Solvent Quality and Solvent Polarity in Polypeptides. Phys. Chem. Chem. Phys. 2023, 25 (6), 4839–4853. 10.1039/d2cp05214h.

(4) Bellissent-Funel, M. C.; Hassanali, A.; Havenith, M.; Henchman, R.; Pohl, P.; Sterpone, F.; Van Der Spoel, D.; Xu, Y.; Garcia, A. E. Water Determines the Structure and Dynamics of Proteins. Chem. Rev. 2016, 116 (13), 7673–7697. 10.1021/acs.chemrev.5b00664.

(5) Pocker, Y. Water in Enzyme Reactions: Biophysical Aspects of Hydration-Dehydration Processes. Cell. Mol. Life Sci. 2000, 57 (7), 1008–1017. 10.1007/pl00000741.

(6) Hassanali, A.; Giberti, F.; Cuny, J.; Kühne, T. D.; Parrinello, M. Proton Transfer through the Water Gossamer. Proc. Natl. Acad. Sci. 2013, 110 (34), 13723–13728. 10.1073/pnas.1306642110.

(7) Li, G.; Wang, B.; Resasco, D. E. Water-Mediated Heterogeneously Catalyzed Reactions. ACS Catal. 2020, 10 (2), 1294–1309. 10.1021/acscatal.9b04637.

(8) Ball, P. Water as an Active Constituent in Cell Biology. Chem. Rev. 2008, 108 (1), 74–108. 10.1021/cr068037a.

(9) Breiten, B.; Lockett, M. R.; Sherman, W.; Fujita, S.; Al-Sayah, M.; Lange, H.; Bowers, C. M.; Heroux, A.; Krilov, G.; Whitesides, G. M. Water Networks Contribute to Enthalpy/Entropy Compensation in Protein–Ligand Binding. J. Am. Chem. Soc. 2013, 135 (41), 15579–15584. 10.1021/ja4075776.

(10) Kokkonen, P.; Sykora, J.; Prokop, Z.; Ghose, A.; Bednar, D.; Amaro, M.; Beerens, K.; Bidmanova, S.; Slanska, M.; Brezovsky, J.; Damborsky, J.; Hof, M. Molecular Gating of an Engineered Enzyme Captured in Real Time. J. Am. Chem. Soc. 2018, 140 (51), 17999– 18008. 10.1021/jacs.8b09848.

(11) Zhu, F.; Tajkhorshid, E.; Schulten, K. Pressure-Induced Water Transport in Membrane Channels Studied by Molecular Dynamics. Biophys. J. 2002, 83 (1), 154–160. 10.1016/S0006-3495(02)75157-6.

(12) Pluhackova, K.; Schittny, V.; Bürkner, P.; Siligan, C.; Horner, A. Multiple Pore Lining Residues Modulate Water Permeability of GlpF. Protein Sci. 2022, 31 (10), e4431. 10.1002/pro.4431.

(13) Jensen, M. Ø.; Mouritsen, O. G. Single-Channel Water Permeabilities of Escherichia Coli Aquaporins AqpZ and GlpF. Biophys. J. 2006, 90 (7), 2270–2284. 10.1529/biophysj.105.073965.

(14) de Groot, B. L.; Grubmüller, H. Water Permeation Across Biological Membranes: Mechanism and Dynamics of Aquaporin-1 and GlpF. Science (80-. ). 2001, 294 (5550), 2353–2357. 10.1126/science.1066115.

(15) Murata, K.; Mitsuoka, K.; Hirai, T.; Walz, T.; Agre, P.; Heymann, J. B.; Engel, A.; Fujiyoshi, Y. Structural Determinants of Water Permeation through Aquaporin-1. Nature 2000, 407 (6804), 599–605. 10.1038/35036519.

(16) Saparov, S. M.; Pohl, P. Beyond the Diffusion Limit: Water Flow through the Empty Bacterial Potassium Channel. Proc. Natl. Acad. Sci. 2004, 101 (14), 4805–4809. 10.1073/pnas.0308309101.

(17) Hadidi, H.; Kamali, R. Molecular Dynamics Study of Water Transport through AQP5-R188C Mutant Causing Palmoplantar Keratoderma (PPK) Using the Gating Mechanism Concept. Biophys. Chem. 2021, 277, 106655. 10.1016/j.bpc.2021.106655.

(18) Beckstein, O.; Biggin, P. C.; Sansom, M. S. P. A Hydrophobic Gating Mechanism for Nanopores. J. Phys. Chem. B 2001, 105 (51), 12902–12905. 10.1021/jp012233y.

(19) Secchi, E.; Marbach, S.; Niguès, A.; Stein, D.; Siria, A.; Bocquet, L. Massive Radius-Dependent Flow Slippage in Carbon Nanotubes. Nature 2016, 537 (7619), 210–213. 10.1038/nature19315.

(20) Xu, J.; Zhu, C.; Wang, Y.; Li, H.; Huang, Y.; Shen, Y.; Francisco, J. S.; Zeng, X. C.; Meng, S. Water Transport through Subnanopores in the Ultimate Size Limit: Mechanism from Molecular Dynamics. Nano Res. 2019, 12 (3), 587–592. 10.1007/s12274-018-2258-7.

(21) Coalson, R. D. Driven Water/Ion Transport through Narrow Nanopores: A Molecular Dynamics Perspective. Faraday Discuss. 2018, 209 (0), 249–257. 10.1039/c8fd00073e.

(22) Horner, A.; Pohl, P. Single-File Transport of Water through Membrane Channels. Faraday Discuss. 2018, 209 (0), 9–33. 10.1039/c8fd00122g.

(23) Pfeffermann, J.; Goessweiner-Mohr, N.; Pohl, P. The Energetic Barrier to Single-File Water Flow through Narrow Channels. Biophysical Reviews. Springer Science and Business Media Deutschland GmbH December 23, 2021, pp 913–923. 10.1007/s12551-021-00875-w.

(24) Li, A.-J.; Nussinov, R. A Set of van Der Waals and Coulombic Radii of Protein Atoms for Molecular and Solvent-Accessible Surface Calculation, Packing Evaluation, and Docking. Proteins Struct. Funct. Genet. 1998, 32 (1), 111–127. 10.1002/(SICI)1097-0134(19980701)32:1<111::AID-PROT12>3.0.CO;2-H.

(25) Gerstein, M.; Chothia, C. Packing at the Protein-Water Interface. Proc. Natl. Acad. Sci. 1996, 93 (19), 10167–10172. 10.1073/pnas.93.19.10167.

(26) Lynch, C. I.; Klesse, G.; Rao, S.; Tucker, S. J.; Sansom, M. S. P. Water Nanoconfined in a Hydrophobic Pore: Molecular Dynamics Simulations of Transmembrane Protein 175 and the Influence of Water Models. ACS Nano 2021, 15 (12), 19098–19108. 10.1021/acsnano.1c06443.

(27) Mitusińska, K.; Bzówka, M.; Magdziarz, T.; Góra, A. Geometry-Based versus Small-Molecule Tracking Method for Tunnel Identification: Benefits and Pitfalls. J. Chem. Inf. Model. 2022, 62 (24), 6803–6811. 10.1021/acs.jcim.2c00985.

(28) Fornage, M.; Lee, C. R.; Doris, P. A.; Bray, M. S.; Heiss, G.; Zeldin, D. C.; Boerwinkle, E. The Soluble Epoxide Hydrolase Gene Harbors Sequence Variation Associated with Susceptibility to and Protection from Incident Ischemic Stroke. Hum. Mol. Genet. 2005, 14 (19), 2829–2837. 10.1093/hmg/ddi315.

(29) Burley, S. K.; Berman, H. M.; Bhikadiya, C.; Bi, C.; Chen, L.; Di Costanzo, L.; Christie, C.; Duarte, J. M.; Dutta, S.; Feng, Z.; Ghosh, S.; Goodsell, D. S.; Green, R. K.; Guranovic, V.; Guzenko, D.; Hudson, B. P.; Liang, Y.; Lowe, R.; Peisach, E.; Periskova, I.; Randle, C.; Rose, A.; Sekharan, M.; Shao, C.; Tao, Y. P.; Valasatava, Y.; Voigt, M.; Westbrook, J.; Young, J.; Zardecki, C.; Zhuravleva, M.; Kurisu, G.; Nakamura, H.; Kengaku, Y.; Cho, H.; Sato, J.; Kim, J. Y.; Ikegawa, Y.; Nakagawa, A.; Yamashita, R.; Kudou, T.; Bekker, G. J.; Suzuki, H.; Iwata, T.; Yokochi, M.; Kobayashi, N.; Fujiwara, T.; Velankar, S.; Kleywegt, G. J.; Anyango, S.; Armstrong, D. R.; Berrisford, J. M.; Conroy, M. J.; Dana, J. M.; Deshpande, M.; Gane, P.; Gáborová, R.; Gupta, D.; Gutmanas, A.; Koča, J.; Mak, L.; Mir, S.; Mukhopadhyay, A.; Nadzirin, N.; Nair, S.; Patwardhan, A.; Paysan-Lafosse, T.; Pravda, L.; Salih, O.; Sehnal, D.; Varadi, M.; Vǎreková, R.; Markley, J. L.; Hoch, J. C.; Romero, P. R.; Baskaran, K.; Maziuk, D.; Ulrich, E. L.; Wedell, J. R.; Yao, H.; Livny, M.; Ioannidis, Y. E. Protein Data Bank: The Single Global Archive for 3D Macromolecular Structure Data. Nucleic Acids Res. 2019, 47 (D1), D520–D528. 10.1093/nar/gky949.

(30) Stepankova, V.; Khabiri, M.; Brezovsky, J.; Pavelka, A.; Sykora, J.; Amaro, M.; Minofar, B.; Prokop, Z.; Hof, M.; Ettrich, R.; Chaloupkova, R.; Damborsky, J. Expansion of Access Tunnels and Active-Site Cavities Influence Activity of Haloalkane Dehalogenases in Organic Cosolvents. ChemBioChem 2013, 14 (7), 890–897. 10.1002/cbic.201200733.

(31) Mowbray, S. L.; Elfström, L. T.; Ahlgren, K. M.; Andersson, C. E.; Widersten, M. X-Ray Structure of Potato Epoxide Hydrolase Sheds Light on Substrate Specificity in Plant Enzymes. Protein Sci. 2006, 15 (7), 1628–1637. 10.1110/ps.051792106.

(32) Grochulski, P.; Li, Y.; Schrag, J. D.; Bouthillier, F.; Smith, P.; Harrison, D.; Rubin, B.; Cygler, M. Insights into Interfacial Activation from an Open Structure of Candida Rugosa Lipase. J. Biol. Chem. 1993, 268 (17), 12843–12847. 10.1016/s0021-9258(18)31464-9.

(33) Grochulski, P.; Li, Y.; Schrag, J. D.; Cygler, M. Two Conformational States of Candida Rugosa Lipase. Protein Sci. 1994, 3 (1), 82–91. 10.1002/pro.5560030111.

(34) Takai, K.; Chiyo, N.; Nakajima, T.; Nariai, T.; Ishikawa, C.; Nakatani, S.; Ikeno, A.; Yamamoto, S.; Sone, T. Three-Dimensional Rational Approach to the Discovery of Potent Substituted Cyclopropyl Urea Soluble Epoxide Hydrolase Inhibitors. Bioorg. Med. Chem. Lett. 2015, 25 (8), 1705–1708. 10.1016/j.bmcl.2015.02.076.

(35) Anandakrishnan, R.; Aguilar, B.; Onufriev, A. V. H++ 3.0: Automating PK Prediction and the Preparation of Biomolecular Structures for Atomistic Molecular Modeling and Simulations. Nucleic Acids Res. 2012, 40 (W1), W537–W541. 10.1093/nar/gks375.

(36) Koudelakova, T.; Bidmanova, S.; Dvorak, P.; Pavelka, A.; Chaloupkova, R.; Prokop, Z.; Damborsky, J. Haloalkane Dehalogenases: Biotechnological Applications. Biotechnol. J. 2013, 8 (1), 32–45. 10.1002/biot.201100486.

(37) Elfström, L. T.; Widersten, M. Catalysis of Potato Epoxide Hydrolase, StEH1. Biochem. J. 2005, 390 (2), 633–640. 10.1042/bj20050526.

(38) Miranda, M.; Urioste, D.; Andrade Souza, L. T.; Mendes, A. A.; De Castro, H. F. Assessment of the Morphological, Biochemical, and Kinetic Properties for Candida Rugosa Lipase Immobilized on Hydrous Niobium Oxide to Be Used in the Biodiesel Synthesis. Enzyme Res. 2011, 2011 (1). 10.4061/2011/216435.

(39) Przybyla-Zawislak, B. D.; Srivastava, P. K.; Vázquez-Matías, J.; Mohrenweiser, H. W.; Maxwell, J. E.; Hammock, B. D.; Bradbury, J. A.; Enayetallah, A. E.; Zeldin, D. C.; Grant, D. F. Polymorphisms in Human Soluble Epoxide Hydrolase. Mol. Pharmacol. 2003, 64 (2), 482–490. 10.1124/mol.64.2.482.

(40) Luchko, T.; Gusarov, S.; Roe, D. R.; Simmerling, C.; Case, D. A.; Tuszynski, J.; Kovalenko, A. Three-Dimensional Molecular Theory of Solvation Coupled with Molecular Dynamics in Amber. J. Chem. Theory Comput. 2010, 6 (3), 607–624. 10.1021/ct900460m.

(41) Sindhikara, D. J.; Yoshida, N.; Hirata, F. Placevent: An Algorithm for Prediction of Explicit Solvent Atom Distribution-Application to HIV-1 Protease and F-ATP Synthase. J. Comput. Chem. 2012, 33 (18), 1536–1543. 10.1002/jcc.22984.

(42) Meyder, A.; Nittinger, E.; Lange, G.; Klein, R.; Rarey, M. Estimating Electron Density Support for Individual Atoms and Molecular Fragments in X-Ray Structures. J. Chem. Inf. Model. 2017, 57 (10), 2437–2447. 10.1021/acs.jcim.7b00391.

(43) Case, D. A.; Ben-Shalom, I. Y.; Brozel, S. R.; Cerutti, D. S.; Cheatham, T. E.; Cruzeiro, V. W. D.; Darden, T. A.; Duke, R. E.; Ghoreishi, D.; Gilson, M. K.; Gohlke, H.; Goetz, A. W.; Greene, D.; Harris, R.; Homeyer, N.; Huang, Y.; Izadi, S.; Kovalenko, A.; Kurtzman, T.; Lee, T. S.; LeGrand, S.; Li, P.; Lin, C.; Liu, J.; Luchko, T.; Luo, R.; Mermelstein, D. J.; Merz, K. M.; Miao, Y.; Monard, G.; Nguyen, C.; Nguyen, H.; Omelyan, I.; Onufriev, A.; Pan, F.; Qi, R.; Roe, D. R.; Roitberg, A.; Sagui, C.; Schott-Verdugo, S.; Shen, J.; Simmerling, C. L.; Smith, J.; Salomon-Ferrer, R.; Swails, J.; Walker, R. C.; Wang, J.; Wei, H.; Wolf, R. M.; Wu, X.; Xiao, L.; York, D. M.; Kollman, P. A. AMBER 18. University of California: San Francisco 2018. ambermd.org.

(44) Izadi, S.; Anandakrishnan, R.; Onufriev, A. V. Building Water Models: A Different Approach. J. Phys. Chem. Lett. 2014, 5 (21), 3863–3871. 10.1021/jz501780a.

(45) Salomon-Ferrer, R.; Götz, A. W.; Poole, D.; Le Grand, S.; Walker, R. C. Routine Microsecond Molecular Dynamics Simulations with AMBER on GPUs. 2. Explicit Solvent Particle Mesh Ewald. J. Chem. Theory Comput. 2013, 9 (9), 3878–3888. 10.1021/ct400314y.

(46) Le Grand, S.; Götz, A. W.; Walker, R. C. SPFP: Speed without Compromise - A Mixed Precision Model for GPU Accelerated Molecular Dynamics Simulations. Comput. Phys. Commun. 2013, 184 (2), 374–380. 10.1016/j.cpc.2012.09.022.

(47) Maier, J. A.; Martinez, C.; Kasavajhala, K.; Wickstrom, L.; Hauser, K. E.; Simmerling, C. Ff14SB: Improving the Accuracy of Protein Side Chain and Backbone Parameters from Ff99SB. J. Chem. Theory Comput. 2015, 11 (8), 3696–3713. 10.1021/acs.jctc.5b00255.

(48) Hopkins, C. W.; Le Grand, S.; Walker, R. C.; Roitberg, A. E. Long-Time-Step Molecular Dynamics through Hydrogen Mass Repartitioning. J. Chem. Theory Comput. 2015, 11 (4), 1864–1874. 10.1021/ct5010406.

(49) Darden, T.; York, D.; Pedersen, L. Particle Mesh Ewald: An N·log(N) Method for Ewald Sums in Large Systems. J. Chem. Phys. 1993, 98 (12), 10089–10092. 10.1063/1.464397.

(50) Essmann, U.; Perera, L.; Berkowitz, M. L.; Darden, T.; Lee, H.; Pedersen, L. G. A Smooth Particle Mesh Ewald Method. J. Chem. Phys. 1995, 103 (19), 8577–8593. 10.1063/1.470117.

(51) Ryckaert, J.-P.; Ciccotti, G.; Berendsen, H. J. . Numerical Integration of the Cartesian Equations of Motion of a System with Constraints: Molecular Dynamics of n-Alkanes. J. Comput. Phys. 1977, 23 (3), 327–341. 10.1016/0021-9991(77)90098-5.

(52) Doerr, S.; Harvey, M. J.; Noé, F.; De Fabritiis, G. HTMD: High-Throughput Molecular Dynamics for Molecular Discovery. J. Chem. Theory Comput. 2016, 12 (4), 1845–1852. 10.1021/acs.jctc.6b00049.

(53) Hruska, E.; Abella, J. R.; Nüske, F.; Kavraki, L. E.; Clementi, C. Quantitative Comparison of Adaptive Sampling Methods for Protein Dynamics. J. Chem. Phys. 2018, 149 (24), 244119. 10.1063/1.5053582.

(54) Betz, R. M.; Dror, R. O. How Effectively Can Adaptive Sampling Methods Capture Spontaneous Ligand Binding? J. Chem. Theory Comput. 2019, 15 (3), 2053–2063. 10.1021/acs.jctc.8b00913.

(55) Pérez-Hernández, G.; Paul, F.; Giorgino, T.; De Fabritiis, G.; Noé, F. Identification of Slow Molecular Order Parameters for Markov Model Construction. J. Chem. Phys. 2013, 139 (1), 015102. 10.1063/1.4811489.

(56) Roe, D. R.; Cheatham, T. E. PTRAJ and CPPTRAJ: Software for Processing and Analysis of Molecular Dynamics Trajectory Data. J. Chem. Theory Comput. 2013, 9 (7), 3084–3095. 10.1021/ct400341p.

(57) Chovancova, E.; Pavelka, A.; Benes, P.; Strnad, O.; Brezovsky, J.; Kozlikova, B.; Gora, A.; Sustr, V.; Klvana, M.; Medek, P.; Biedermannova, L.; Sochor, J.; Damborsky, J. CAVER 3.0: A Tool for the Analysis of Transport Pathways in Dynamic Protein Structures. PLoS Comput. Biol. 2012, 8 (10), e1002708. 10.1371/journal.pcbi.1002708.

(58) Magdziarz, T.; Mitusińska, K.; Bzówka, M.; Raczyńska, A.; Stańczak, A.; Banas, M.; Bagrowska, W.; Góra, A. AQUA-DUCT 1.0: Structural and Functional Analysis of Macromolecules from an Intramolecular Voids Perspective. Bioinformatics 2020, 36 (8), 2599–2601. 10.1093/bioinformatics/btz946.

(59) Brezovsky, J.; Thirunavukarasu, A. S.; Surpeta, B.; Sequeiros-Borja, C. E.; Mandal, N.; Sarkar, D. K.; Dongmo Foumthuim, C. J.; Agrawal, N. TransportTools: A Library for High-Throughput Analyses of Internal Voids in Biomolecules and Ligand Transport through Them. Bioinformatics 2022, 38 (6), 1752–1753. 10.1093/bioinformatics/btab872.

(60) Sequeiros-Borja, C.; Surpeta, B.; Marchlewski, I.; Brezovsky, J. Divide-and-Conquer Approach to Study Protein Tunnels in Long Molecular Dynamics Simulations. MethodsX 2023, 10, 101968. 10.1016/j.mex.2022.101968.

(61) Petřek, M.; Otyepka, M.; Banáš, P.; Košinová, P.; Koča, J.; Damborský, J. CAVER: A New Tool to Explore Routes from Protein Clefts, Pockets and Cavities. BMC Bioinformatics 2006, 7 (1), 316. 10.1186/1471-2105-7-316.

(62) Klvana, M.; Pavlova, M.; Koudelakova, T.; Chaloupkova, R.; Dvorak, P.; Prokop, Z.; Stsiapanava, A.; Kuty, M.; Kuta-Smatanova, I.; Dohnalek, J.; Kulhanek, P.; Wade, R. C.; Damborsky, J. Pathways and Mechanisms for Product Release in the Engineered Haloalkane Dehalogenases Explored Using Classical and Random Acceleration Molecular Dynamics Simulations. J. Mol. Biol. 2009, 392 (5), 1339–1356. 10.1016/j.jmb.2009.06.076.

(63) Mitusińska, K.; Magdziarz, T.; Bzówka, M.; Stańczak, A.; Gora, A. Exploring Solanum Tuberosum Epoxide Hydrolase Internal Architecture by Water Molecules Tracking. Biomolecules 2018, 8 (4), 143. 10.3390/biom8040143.

(64) Rodríguez-Salarichs, J.; García De Lacoba, M.; Prieto, A.; Martínez, M. J.; Barriuso, J. Versatile Lipases from the Candida Rugosa-like Family: A Mechanistic Insight Using Computational Approaches. J. Chem. Inf. Model. 2021, 61 (2), 913–920. 10.1021/acs.jcim.0c01151.

(65) Mancheño, J. M.; Pernas, M. A.; Martínez, M. J.; Ochoa, B.; Rúa, M. L.; Hermoso, J. A. Structural Insights into the Lipase/Esterase Behavior in the Candida Rugosa Lipases Family: Crystal Structure of the Lipase 2 Isoenzyme at 1.97Å Resolution. J. Mol. Biol. 2003, 332 (5), 1059–1069. 10.1016/j.jmb.2003.08.005.

(66) Domínguez de María, P.; Sánchez-Montero, J. M.; Sinisterra, J. V.; Alcántara, A. R. Understanding Candida Rugosa Lipases: An Overview. Biotechnol. Adv. 2006, 24 (2), 180–196. 10.1016/j.biotechadv.2005.09.003.

(67) de Melo, J. J. C.; Gonçalves, J. R.; Brandão, L. M. de S.; Souza, R. L.; Pereira, M. M.; Lima, Á. S.; Soares, C. M. F. Evaluation of Lipase Access Tunnels and Analysis of Substance Transport in Comparison with Experimental Data. Bioprocess Biosyst. Eng. 2022, 45 (7), 1149–1162. 10.1007/s00449-022-02731-x.

(68) Foresti, M. L.; Ferreira, M. L. Computational Approach to Solvent-Free Synthesis of Ethyl Oleate Using Candida r Ugosa and Candida a Ntarctica B Lipases. I. Interfacial Activation and Substrate (Ethanol, Oleic Acid) Adsorption. Biomacromolecules 2004, 5 (6), 2366–2375. 10.1021/bm049688o.

(69) Bzówka, M.; Mitusińska, K.; Hopko, K.; Góra, A. Computational Insights into the Known Inhibitors of Human Soluble Epoxide Hydrolase. Drug Discov. Today 2021, 26 (8), 1914–1921. 10.1016/j.drudis.2021.05.017.

(70) Mitusińska, K.; Wojsa, P.; Bzówka, M.; Raczyńska, A.; Bagrowska, W.; Samol, A.; Kapica, P.; Góra, A. Structure-Function Relationship between Soluble Epoxide Hydrolases Structure and Their Tunnel Network. Comput. Struct. Biotechnol. J. 2022, 20, 193–205. 10.1016/j.csbj.2021.10.042.

(71) McInnes, L.; Healy, J.; Astels, S. Hdbscan: Hierarchical Density Based Clustering. J. Open Source Softw. 2017, 2 (11), 205. 10.21105/joss.00205.

(72) Kajander, T.; Kahn, P. C.; Passila, S. H.; Cohen, D. C.; Lehtiö, L.; Adolfsen, W.; Warwicker, J.; Schell, U.; Goldman, A. Buried Charged Surface in Proteins. Structure 2000, 8 (11), 1203–1214. 10.1016/S0969-2126(00)00520-7.

(73) Brezovsky, J.; Chovancova, E.; Gora, A.; Pavelka, A.; Biedermannova, L.; Damborsky, J. Software Tools for Identification, Visualization and Analysis of Protein Tunnels and Channels. Biotechnol. Adv. 2013, 31 (1), 38–49. 10.1016/j.biotechadv.2012.02.002.

(74) Pavelka, A.; Sebestova, E.; Kozlikova, B.; Brezovsky, J.; Sochor, J.; Damborsky, J. CAVER: Algorithms for Analyzing Dynamics of Tunnels in Macromolecules. IEEE/ACM Trans. Comput. Biol. Bioinforma. 2016, 13 (3), 505–517. 10.1109/tcbb.2015.2459680.

(75) Bzówka, M.; Mitusińska, K.; Raczyńska, A.; Skalski, T.; Samol, A.; Bagrowska, W.; Magdziarz, T.; Góra, A. Evolution of Tunnels in α/β-Hydrolase Fold Proteins—What Can We Learn from Studying Epoxide Hydrolases? PLOS Comput. Biol. 2022, 18 (5), e1010119. 10.1371/journal.pcbi.1010119.

(76) Schmitt, J.; Brocca, S.; Schmid, R. D.; Pleiss, J. Blocking the Tunnel: Engineering of Candida Rugosa Lipase Mutants with Short Chain Length Specificity. Protein Eng. Des. Sel. 2002, 15 (7), 595–601. 10.1093/protein/15.7.595.

(77) Wang, L.; Althoff, E. A.; Bolduc, J.; Jiang, L.; Moody, J.; Lassila, J. K.; Giger, L.; Hilvert, D.; Stoddard, B.; Baker, D. Structural Analyses of Covalent Enzyme-Substrate Analog Complexes Reveal Strengths and Limitations of de Novo Enzyme Design. J. Mol. Biol. 2012, 415 (3), 615–625. 10.1016/j.jmb.2011.10.043.

(78) Wagner, K.; Vito, S.; Inceoglu, B.; Hammock, B. D. The Role of Long Chain Fatty Acids and Their Epoxide Metabolites in Nociceptive Signaling. Prostaglandins Other Lipid Mediat. 2014, *113–115*, 2–12. 10.1016/j.prostaglandins.2014.09.001.

(79) Decker, M.; Adamska, M.; Cronin, A.; Di Giallonardo, F.; Burgener, J.; Marowsky, A.; Falck, J. R.; Morisseau, C.; Hammock, B. D.; Gruzdev, A.; Zeldin, D. C.; Arand, M. EH3 (ABHD9): The First Member of a New Epoxide Hydrolase Family with High Activity for Fatty Acid Epoxides. J. Lipid Res. 2012, 53 (10), 2038–2045. 10.1194/jlr.m024448.

(80) Morisseau, C.; Hammock, B. D. Impact of Soluble Epoxide Hydrolase and Epoxyeicosanoids on Human Health. Annu. Rev. Pharmacol. Toxicol. 2013, 53 (1), 37–58. 10.1146/annurev-pharmtox-011112-140244.

(81) Kadaoluwa Pathirannahalage, S. P.; Meftahi, N.; Elbourne, A.; Weiss, A. C. G.; McConville, C. F.; Padua, A.; Winkler, D. A.; Costa Gomes, M.; Greaves, T. L.; Le, T. C.; Besford, Q. A.; Christofferson, A. J. Systematic Comparison of the Structural and Dynamic Properties of Commonly Used Water Models for Molecular Dynamics Simulations. J. Chem. Inf. Model. 2021, 61 (9), 4521–4536. 10.1021/acs.jcim.1c00794.

(82) Agrawal, N.; Brezovsky, J. Impact of Water Models on Structure and Dynamics of Ligand-Transport Tunnels in Enzymes Derived from Molecular Dynamics Simulations. bioRxiv 2023, 2023.04.19.537534. 10.1101/2023.04.19.537534.

(83) Efron, B. The Jackknife, the Bootstrap and Other Resampling Plans. The Jackknife, the Bootstrap and Other Resampling Plans 1982. 10.1137/1.9781611970319.

(84) Brezovsky, J.; Babkova, P.; Degtjarik, O.; Fortova, A.; Gora, A.; Iermak, I.; Rezacova, P.; Dvorak, P.; Smatanova, I. K.; Prokop, Z.; Chaloupkova, R.; Damborsky, J. Engineering a de Novo Transport Tunnel. ACS Catal. 2016, 6 (11), 7597–7610. 10.1021/acscatal.6b02081.

