## Supporting information for "Water will find its way: transport through narrow tunnels in hydrolases"

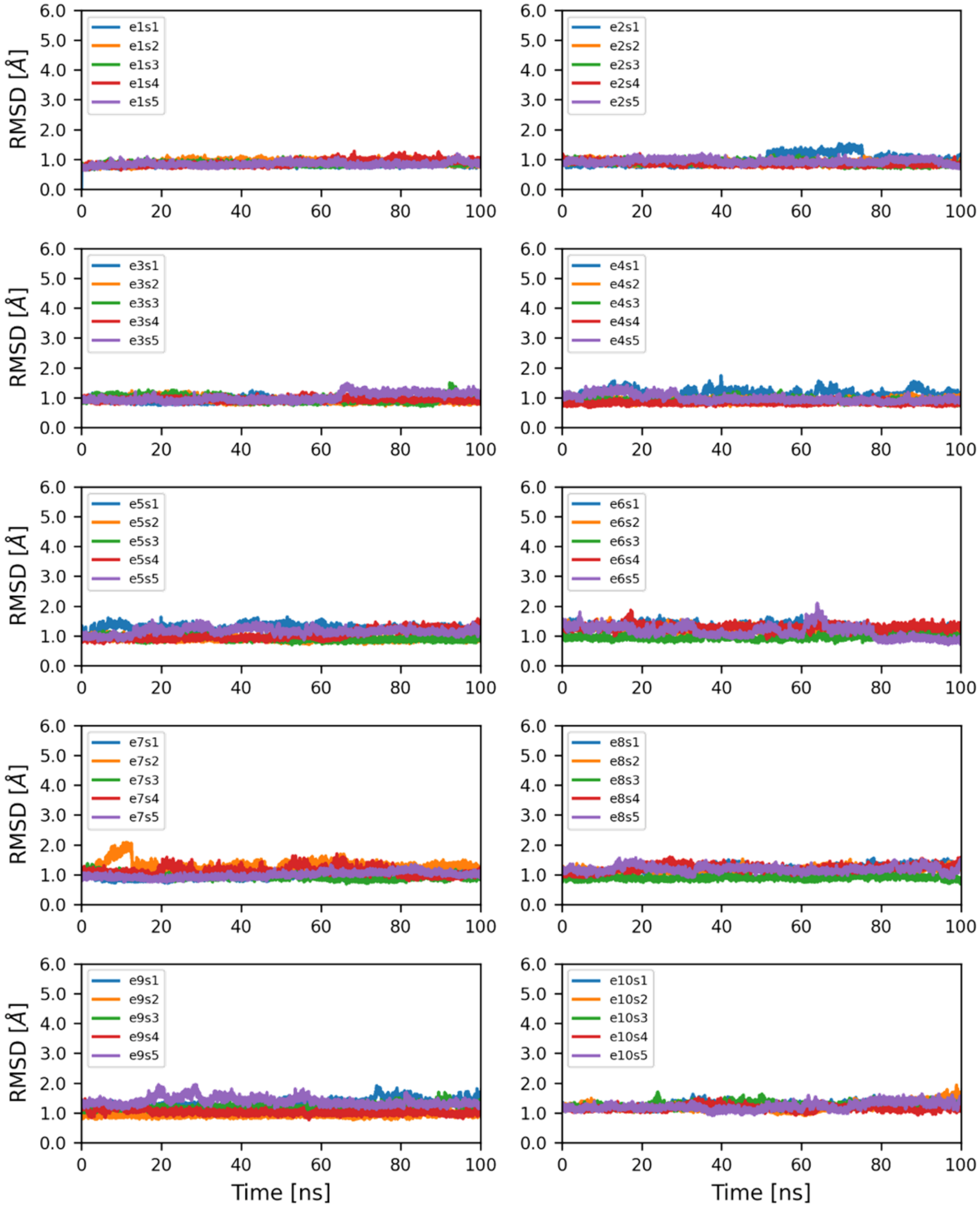


**Figure S1. RMSD of Hal in adaptive MD simulations**. 50 MD simulations of Hal grouped by epochs.

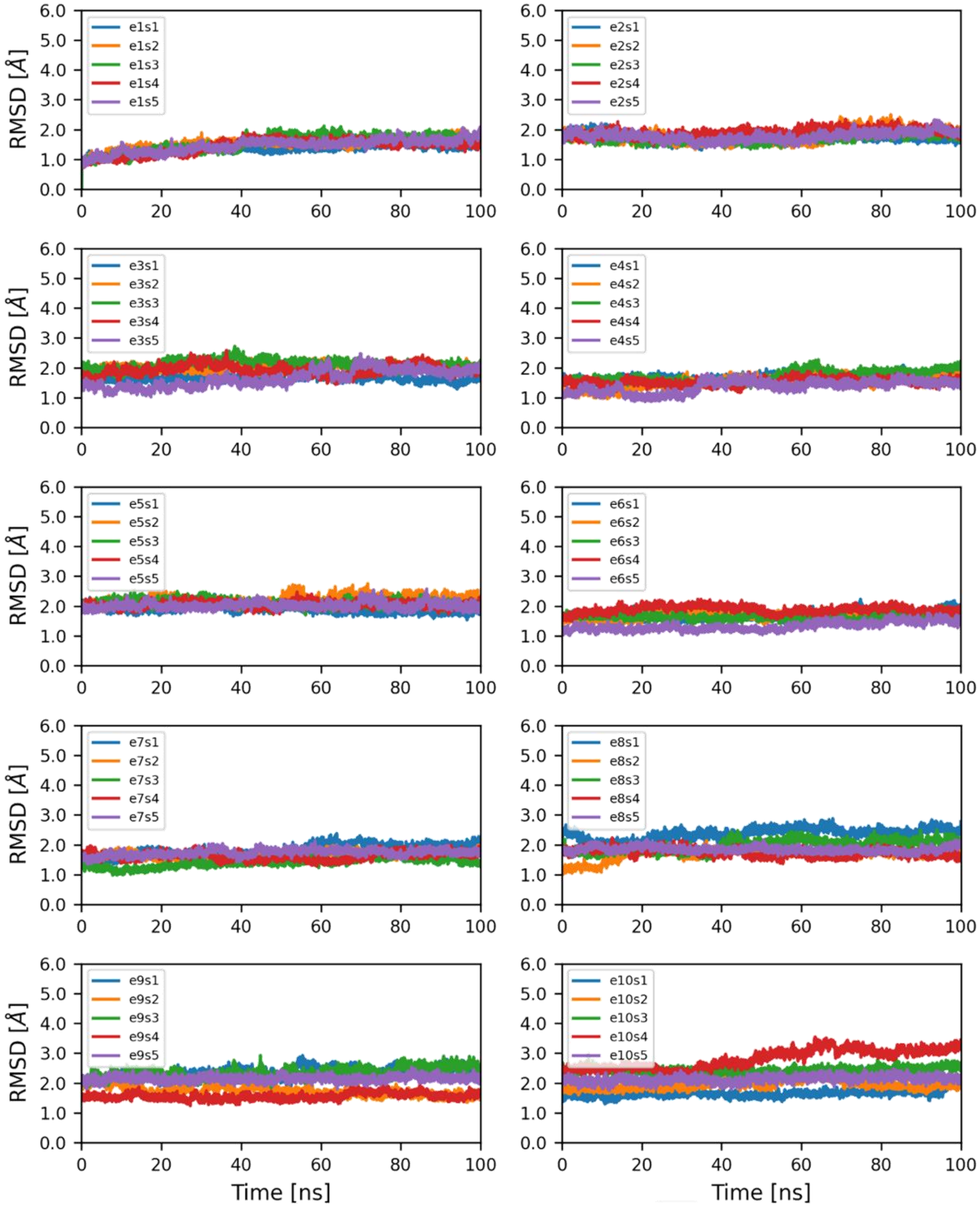


**Figure S2. RMSD of Epx in adaptive MD simulations**. 50 MD simulations of Epx grouped by epochs.

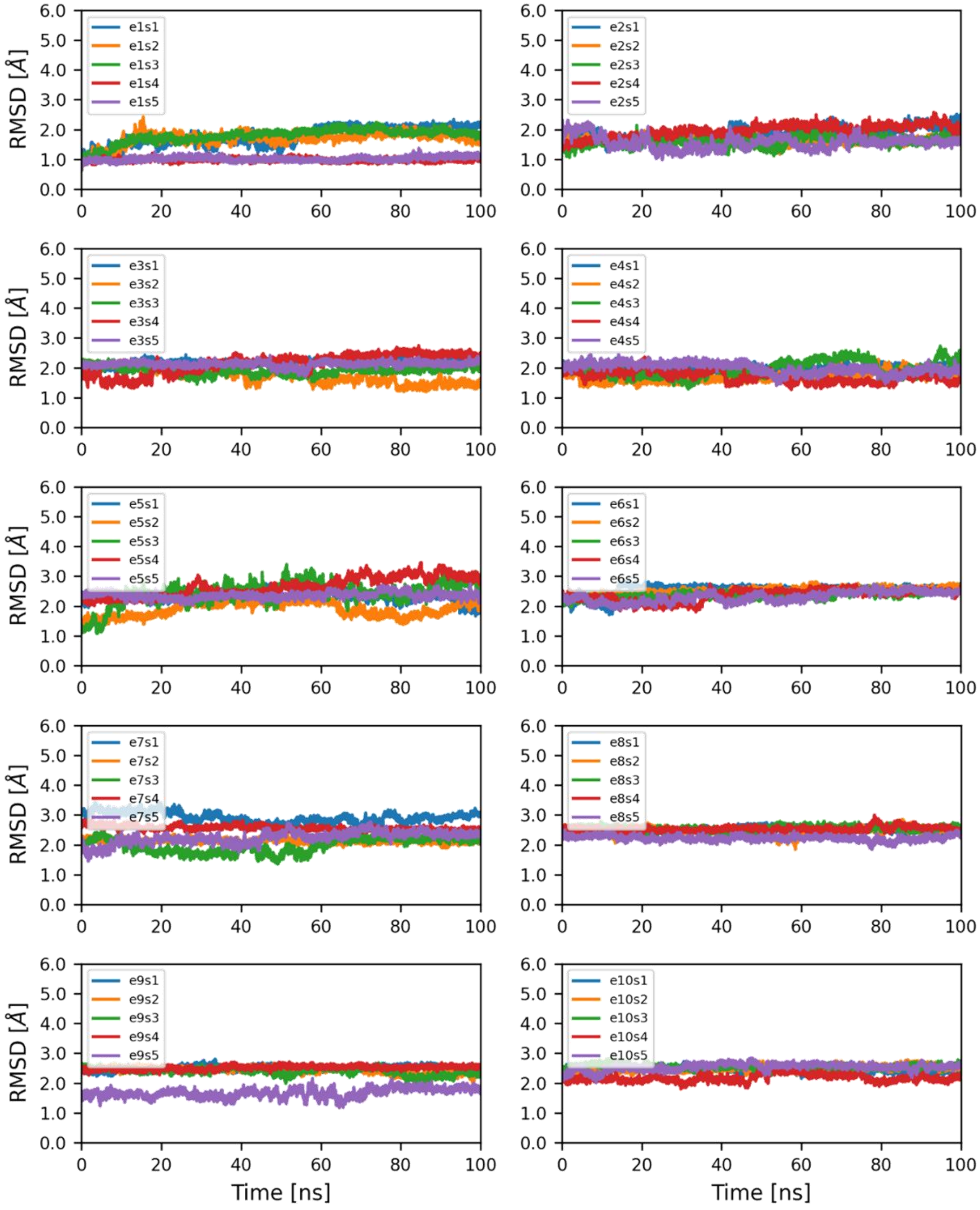


**Figure S3. RMSD of Lip in adaptive MD simulations**. 50 MD simulations of Lip grouped by epochs.

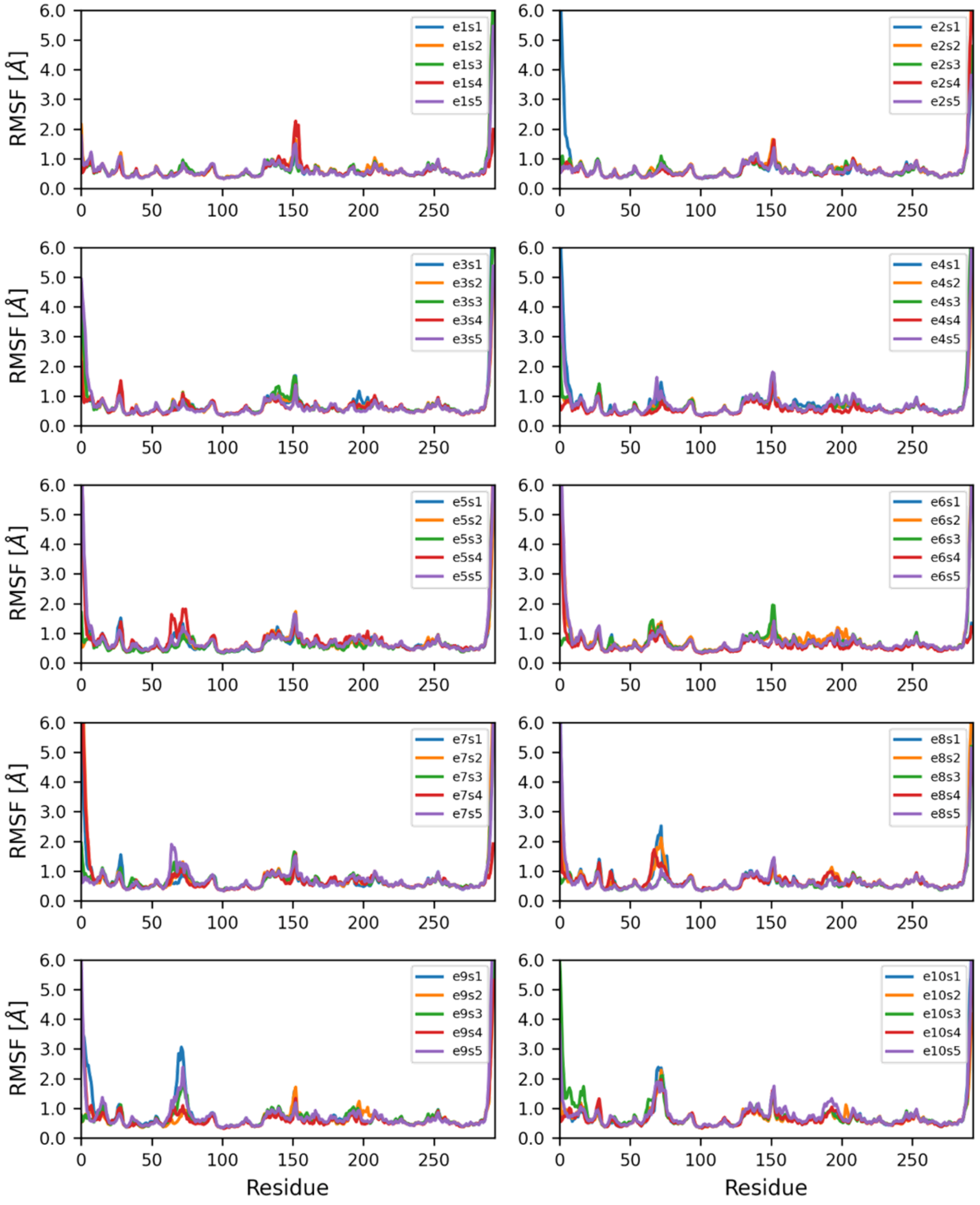


**Figure S4. RMSF of Hal in adaptive MD simulations**. 50 MD simulations of Hal grouped by epochs.

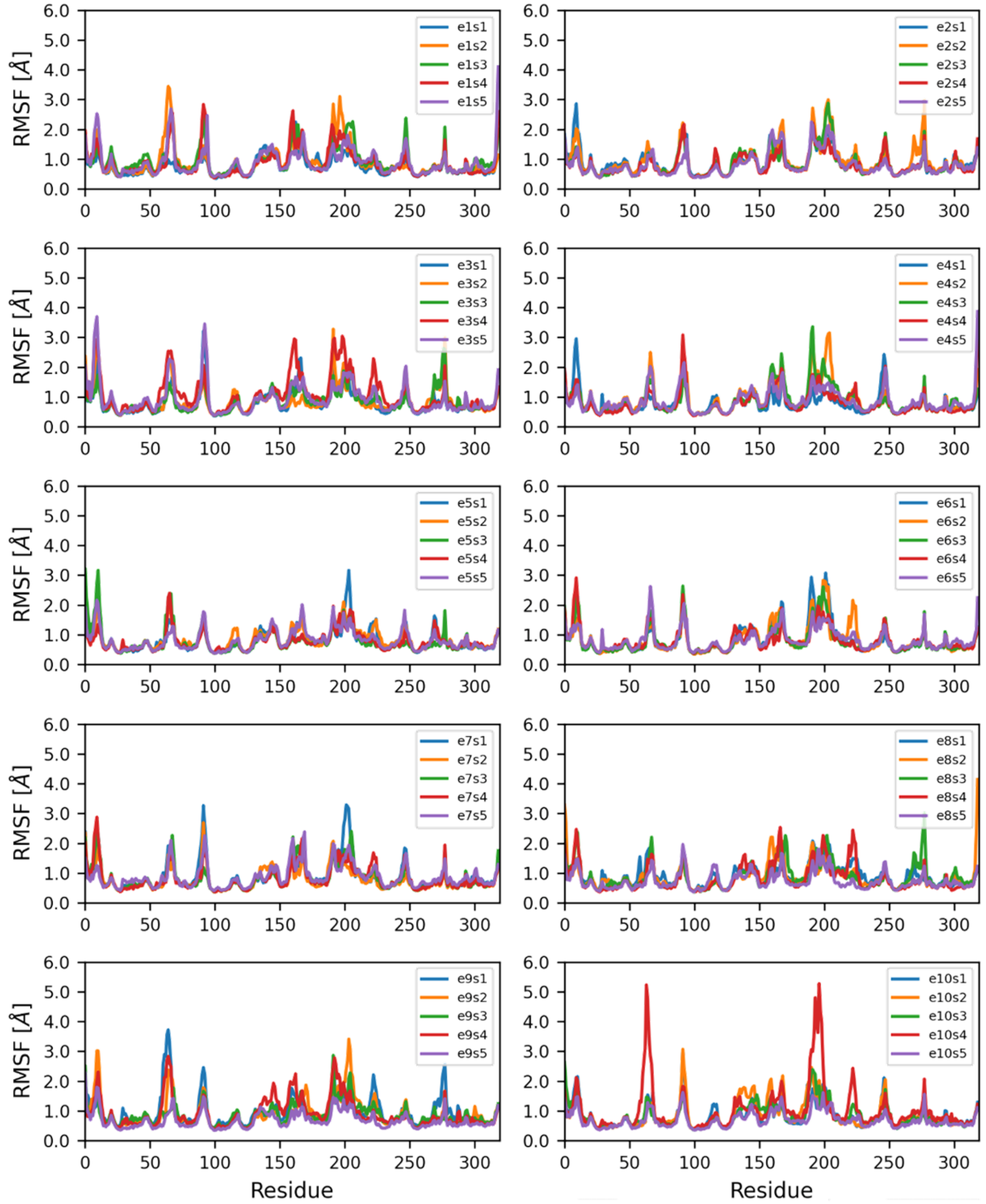


**Figure S5. RMSF of Epx in adaptive MD simulations**. 50 MD simulations of Epx grouped by epochs.

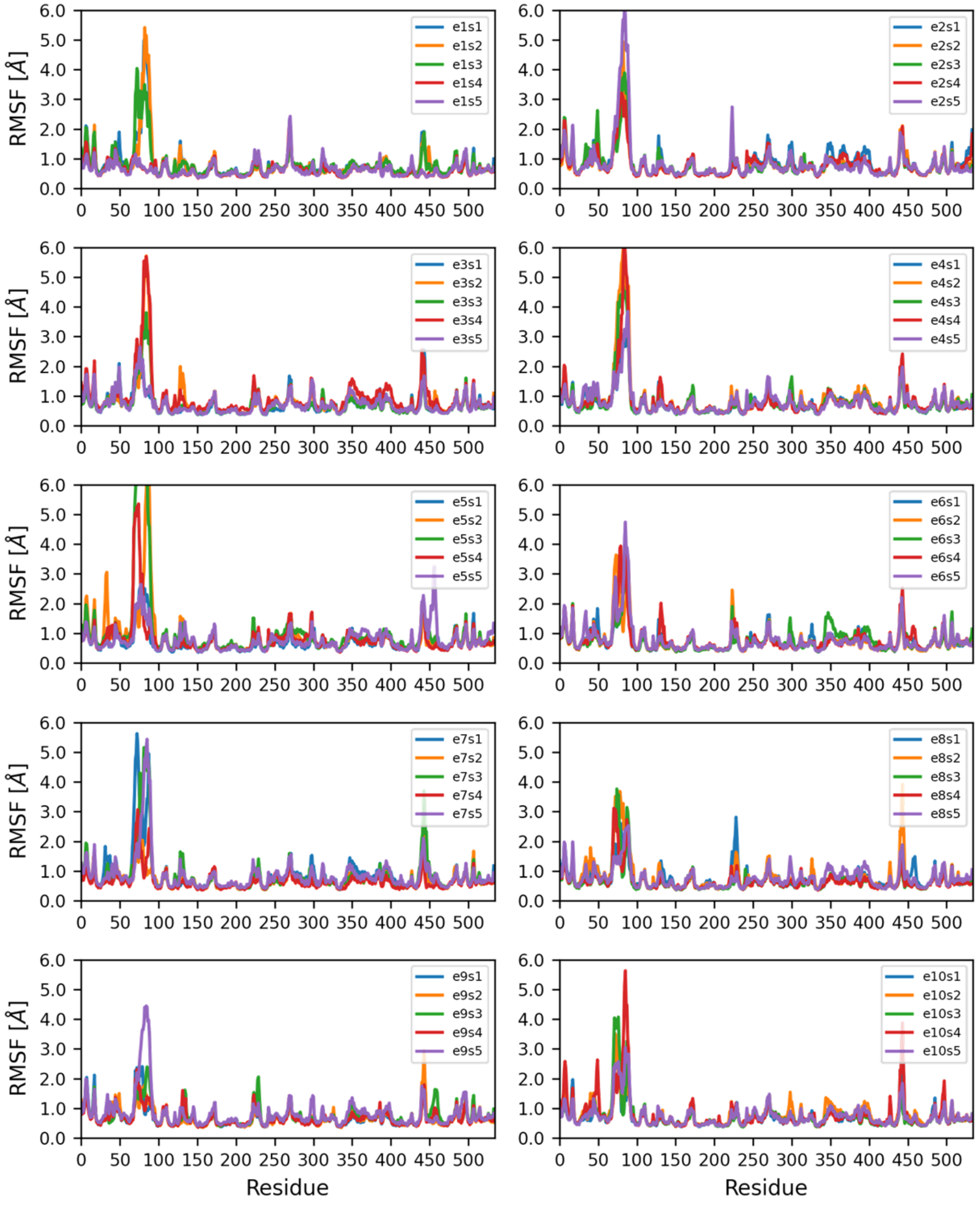


**Figure S6. RMSF of Lip in adaptive MD simulations**. 50 MD simulations of Lip grouped by epochs.

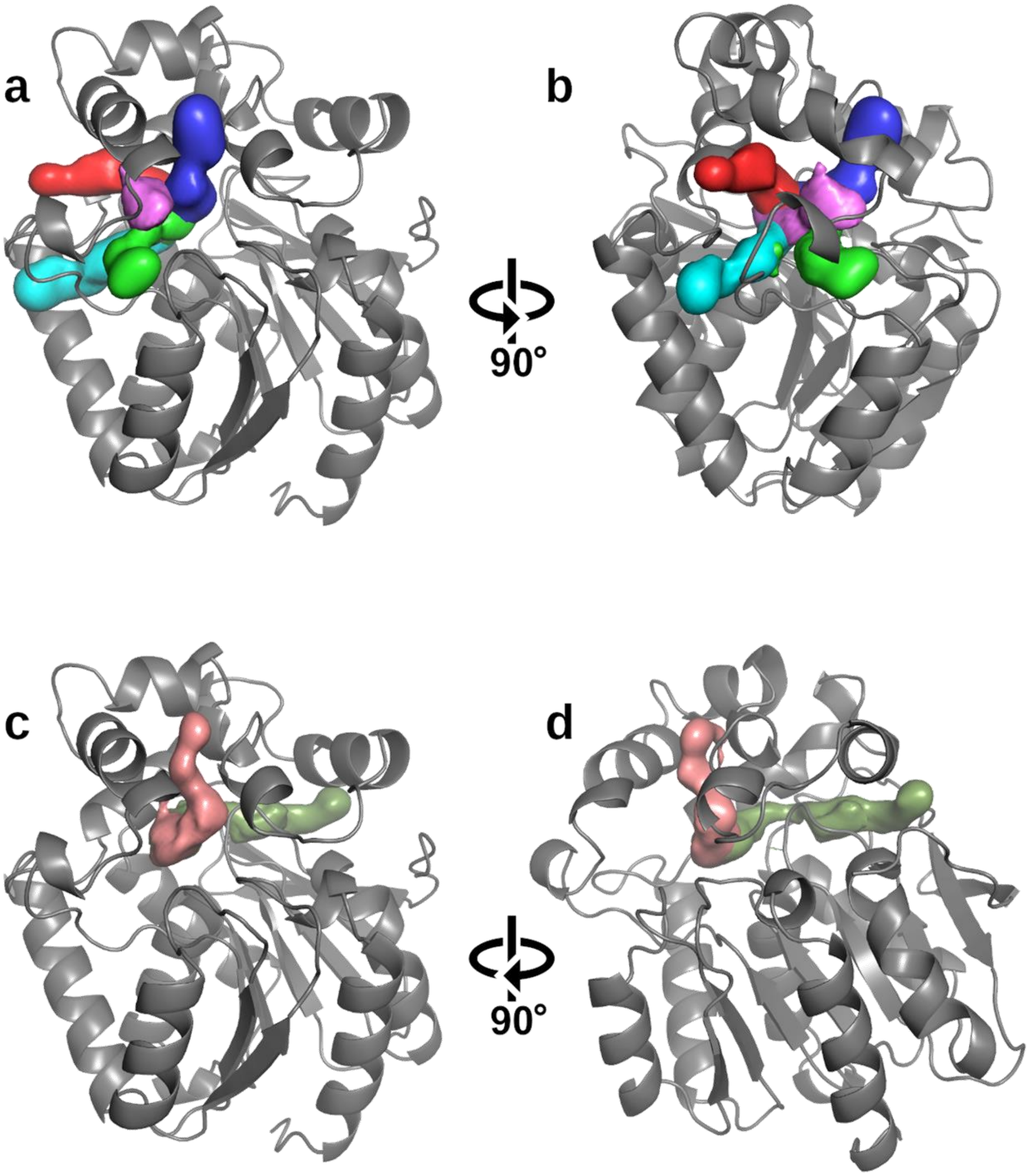


**Figure S7. Tunnels identified in Hal**. Tunnels' densities at 20% of presence in Hal obtained from 5 µs adaptive MD simulations. Tunnels described in the literature: a) *p1* (blue), *p2a* (green), *p2b* (pink), *p2c* (cyan), and *p3* (red); b) rotating the same view 90° counterclockwise in the Z-axis; and newly observed tunnels: c) *up* (salmon) and *side* (light green) tunnels; d) rotating the same view 90° clockwise in the Z-axis.

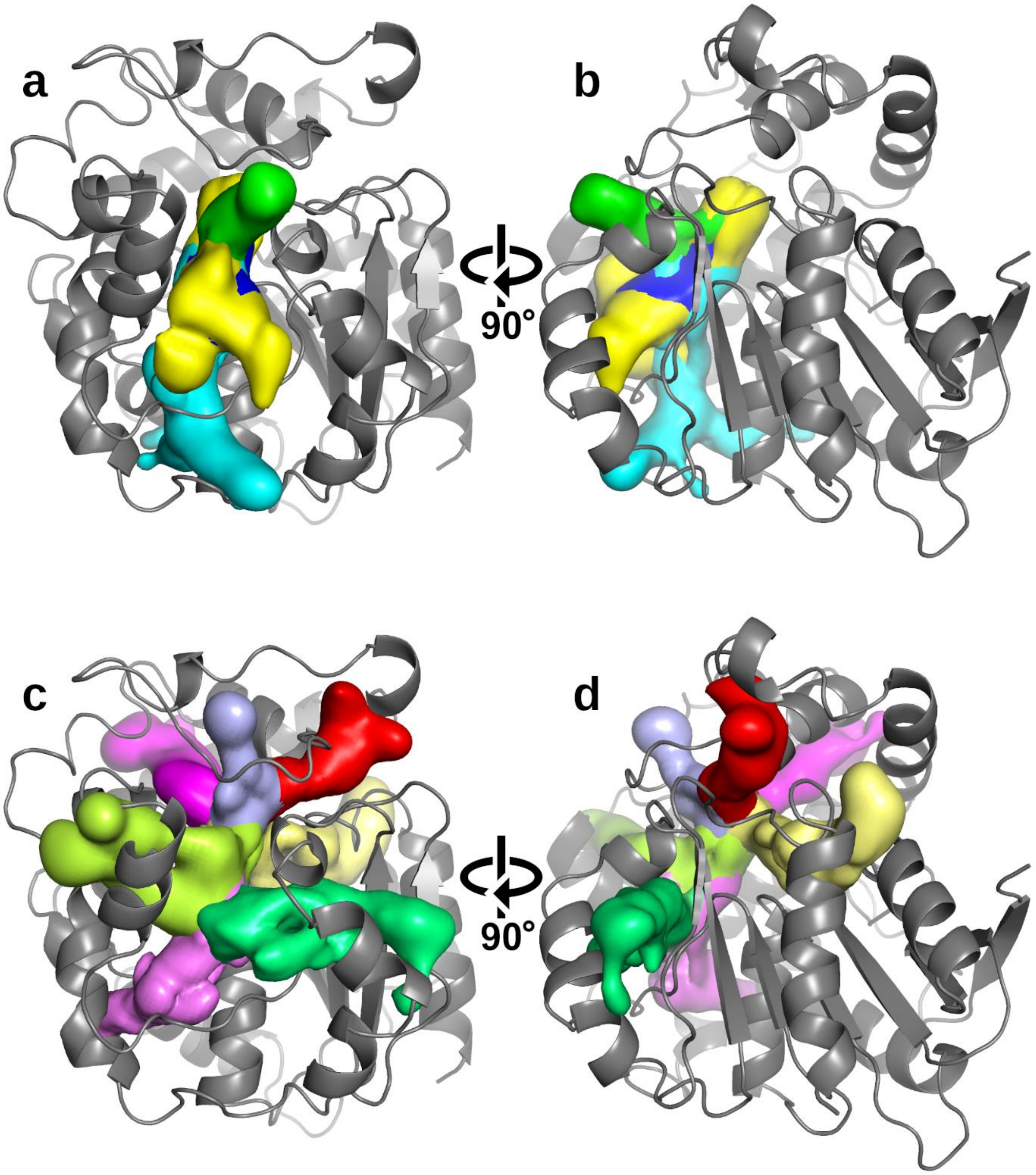


**Figure S8.** T**unnels identified in Epx.** Tunnels' densities at 20% of presence in Epx obtained from 5 µs adaptive MD simulations. Tunnels described in the literature: a) *TM1* (blue and yellow), *TC/M* (green), and *TM2* (cyan); b) rotating the same view 90° clockwise in the Z-axis. Newly identified lesser tunnels that also participate in water transport: c) and the same view d) after rotating the view 90° clockwise in the Z-axis (4, 6, 8, 14, 15, 16, and 21 in red, magenta, light-blue, lemon, pale-yellow, violet, and lime respectively; IDs from **Table S1**).

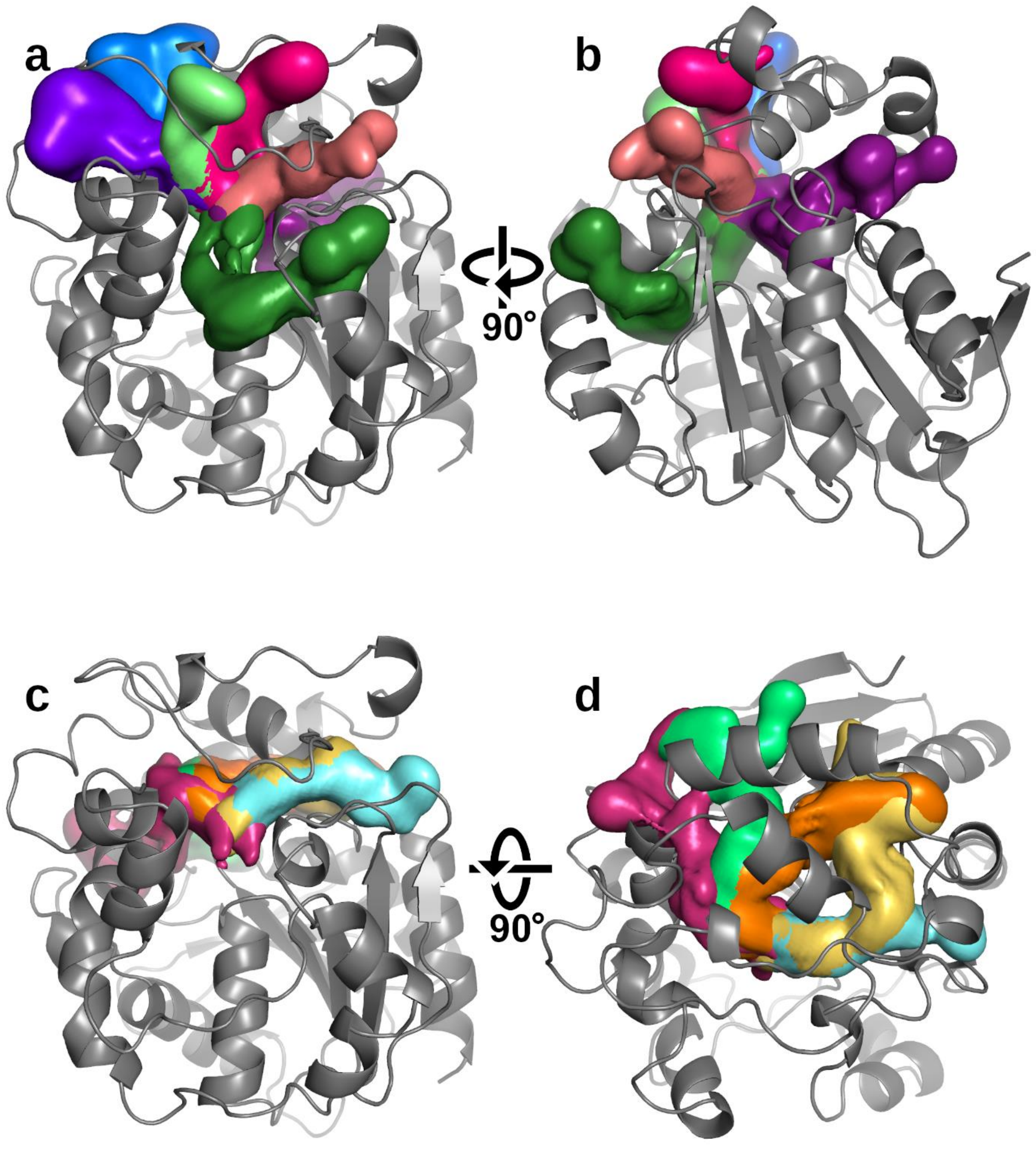


**Figure S9. New water-conducting tunnels identified in Epx.** Tunnels' densities at 20% of presence in Epx obtained from 5 µs adaptive MD simulations. a) and same view b) after rotating the view 90° clockwise in the Z-axis (9, 10, 11, 12, 13, 22, and 25 in blue, light-green, salmon, fuchsia, purple, dark-green, and wine, respectively; IDs from **Table S1**). More tunnels identified c) and the same view d) after rotating the view 90° on the X-axis (7, 17, 19, 23, and 24 in lime, orange, aquamarine, dark-pink, and pale-yellow, respectively; IDs from **Table S1**).

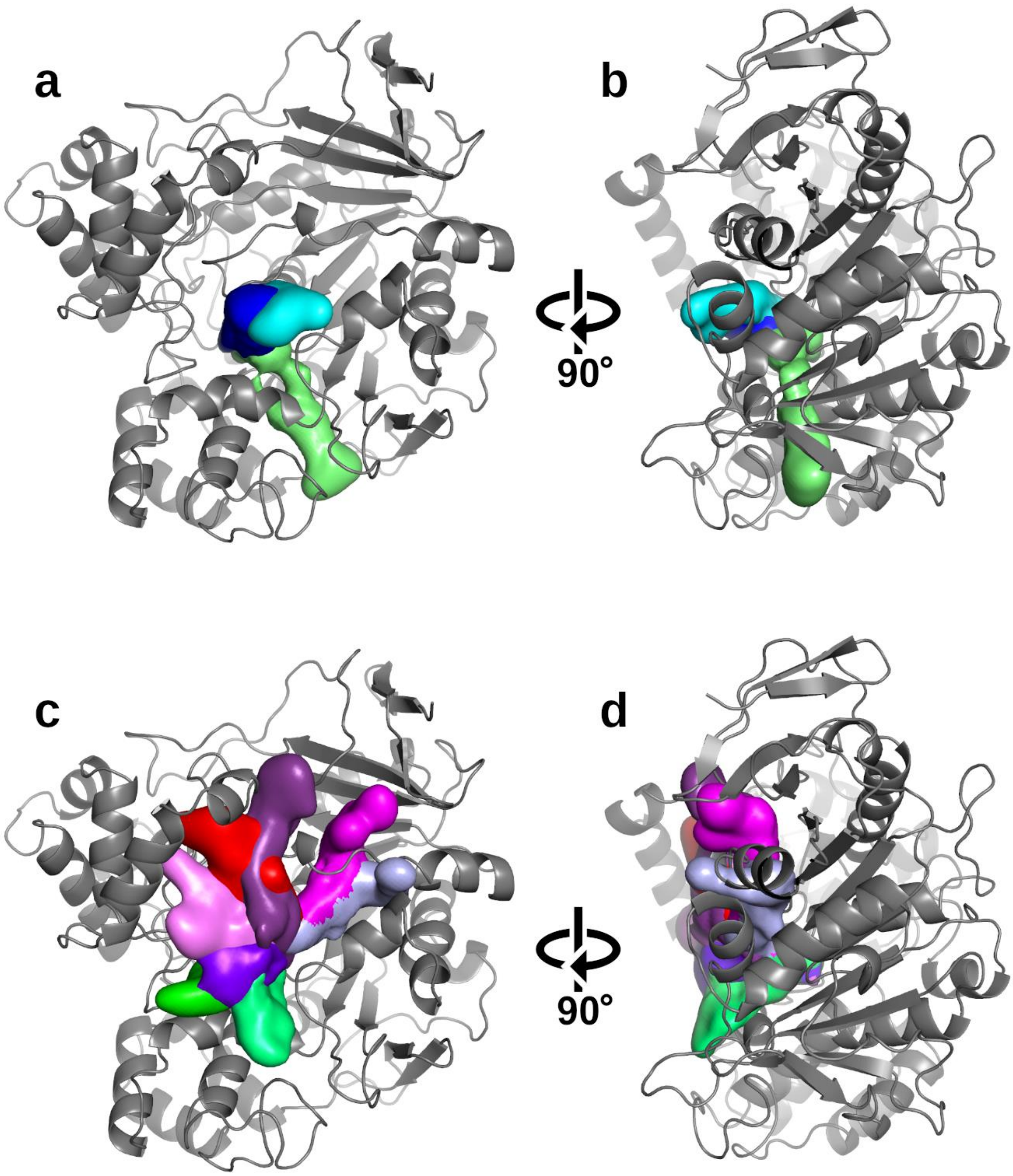


**Figure S10. Tunnels identified in Lip.** Tunnels' densities at 20% of presence in Lip obtained from 5 µs adaptive MD simulations. Tunnels described in the literature: a) the *main* tunnel (blue and cyan) and *ester* tunnel (pale green); b) rotating the same view 90° clockwise in the Z-axis. Newly identified lesser tunnels that also participate in water transport: c) and the same view d) after rotating the view 90° clockwise in the Z-axis (2, 3, 5, 8, 11, 12, 14, and 16 in red, green, magenta, light-blue, lime, purple, dark-purple, and pink, respectively; IDs from **Table S1**).

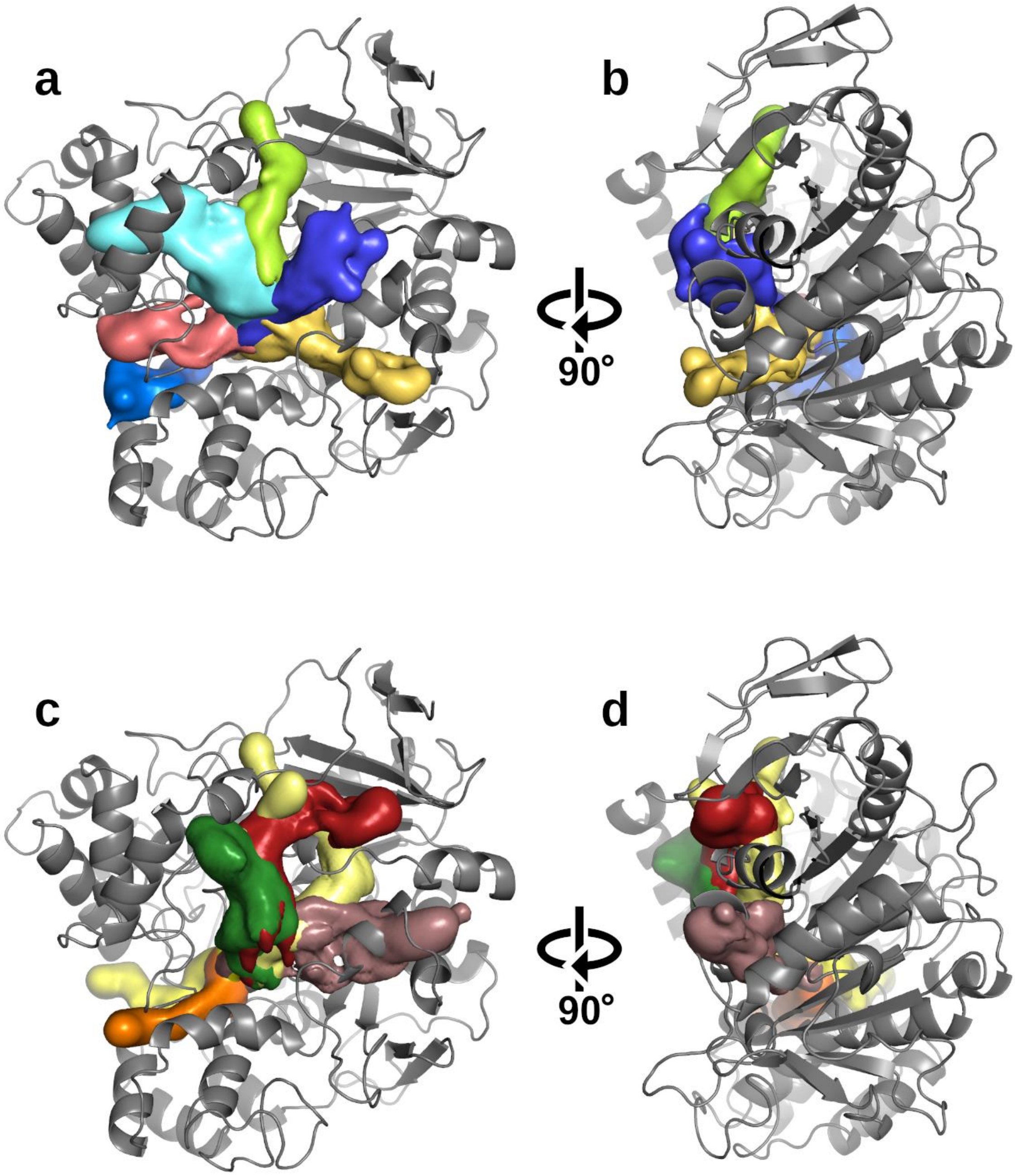


**Figure S11. New water-conducting tunnels identified in Lip**. Tunnels' densities at 20% of presence in Lip obtained from 5 µs adaptive MD simulations. a) and same view b) after rotating the view 90° clockwise in the Z-axis (9, 10, 15, 18, 24, and 26 in salmon, blue, lemon, cyan, pale-yellow, and dark-blue, respectively; IDs from **Table S1**). More tunnels identified c) and the same view d) after rotating the view 90° on the X-axis (6, 17, 19, 38, 66, and 70 in yellow, pale-yellow, orange, dark-red, dark-green, and pale-violet, respectively; IDs from **Table S1**).

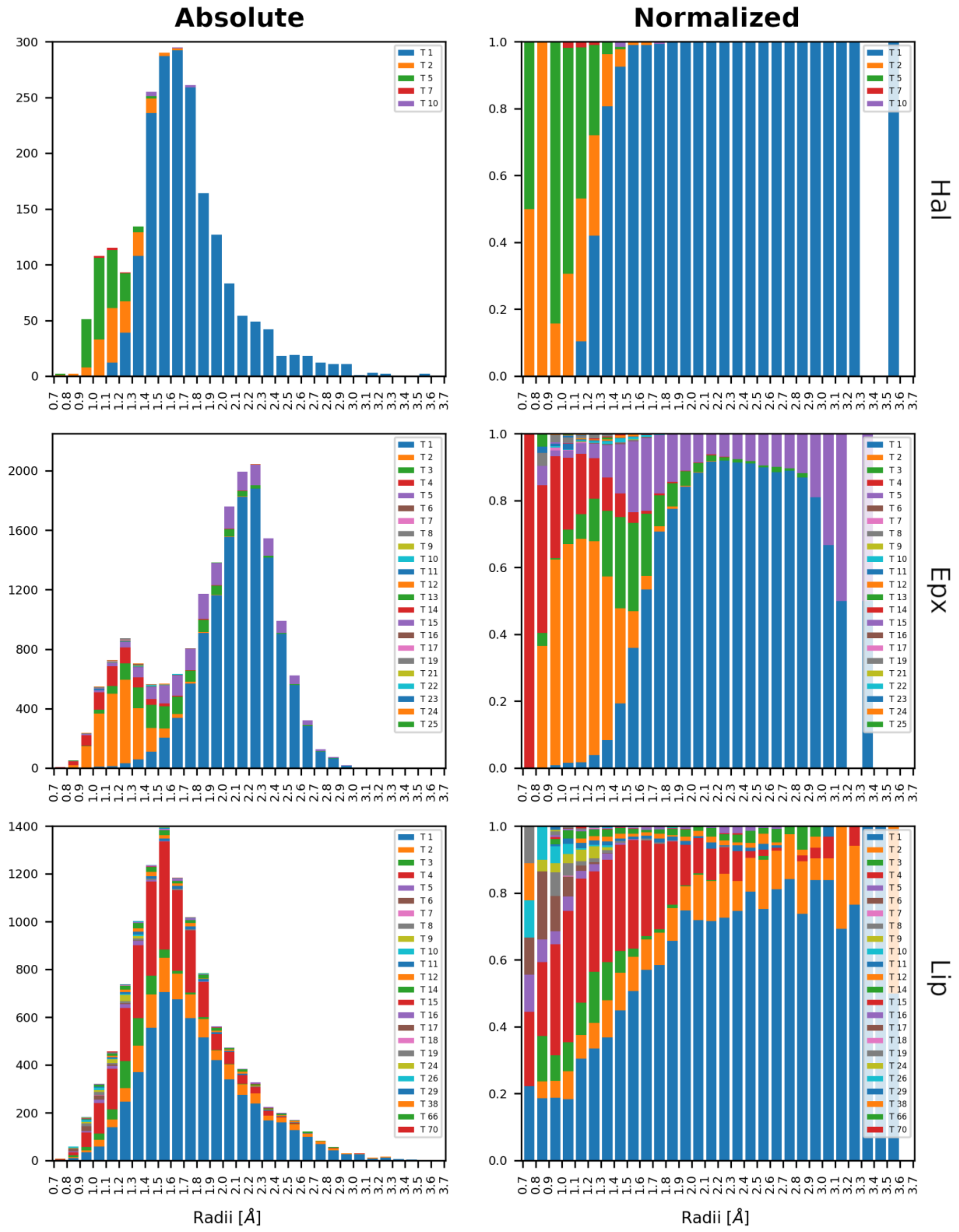


**Figure S12. Distribution of *minimal event sphere* radii in Hal, Epx, and Lip by tunnels**. Histograms of the radius at which a water event occurs are presented in absolute and normalized scales for Hal, Epx, and Lip. Colors represent individual clusters. However, it loops in Epx and Lip cases due to the high number of tunnels identified, IDs from **Table S1**.

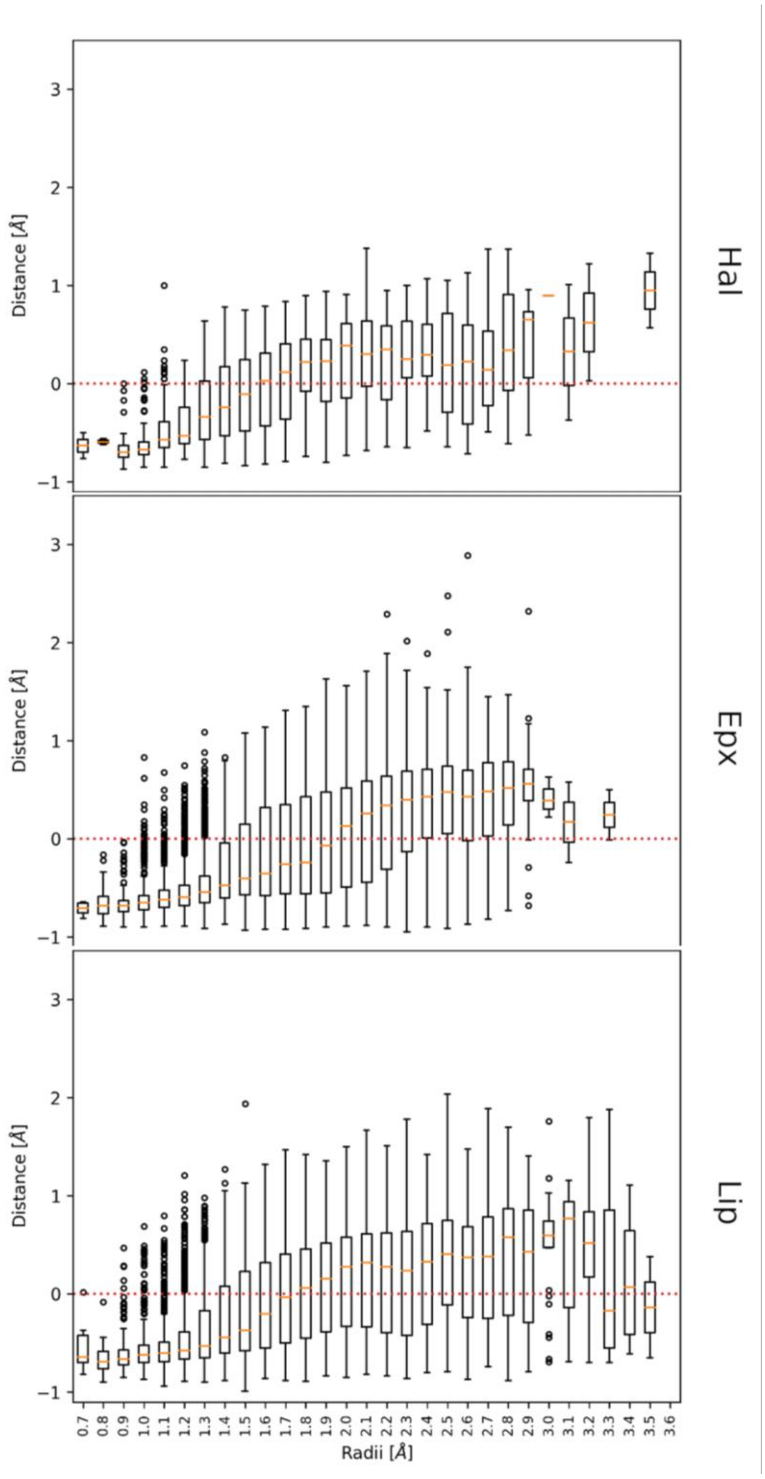


**Figure S13. The surface-surface distances between closest protein atoms and water molecules present in individual bins of *minimal event sphere* radii in Hal, Epx, and Lip**. These distances are calculated as distance between atom centers minus a sum of atomic radii. Hence, distances below the 0 Å threshold (red dotted line) represent the overlap between the closest protein and water atoms.

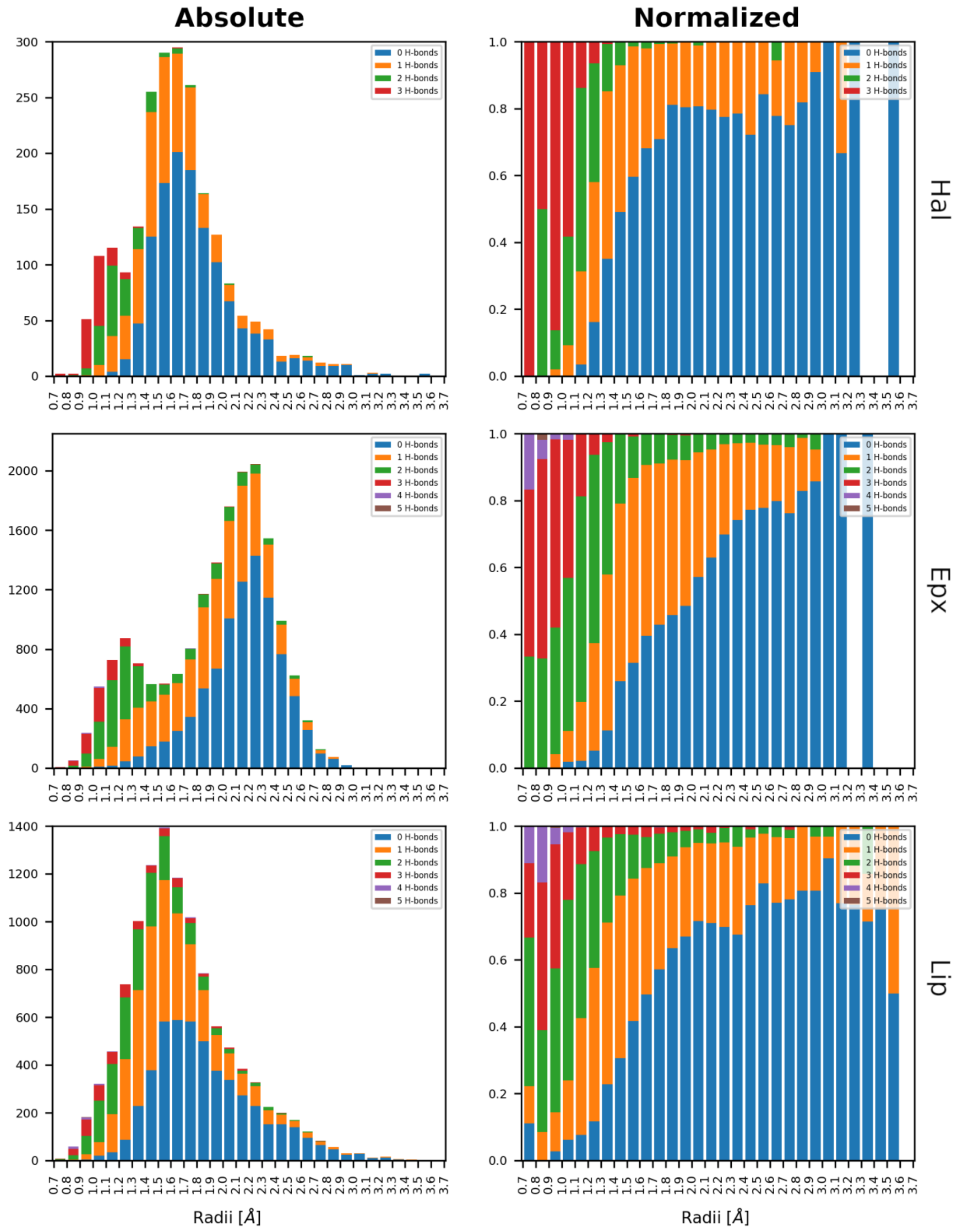


**Figure S14. Distribution of *minimal event sphere* radii in Hal, Epx, and Lip by H-bonds.** Histograms of the radius at which a water event occurs are presented in absolute and normalized scales for Hal, Epx, and Lip. Colors represent the number of H-bonds present in each transport event.

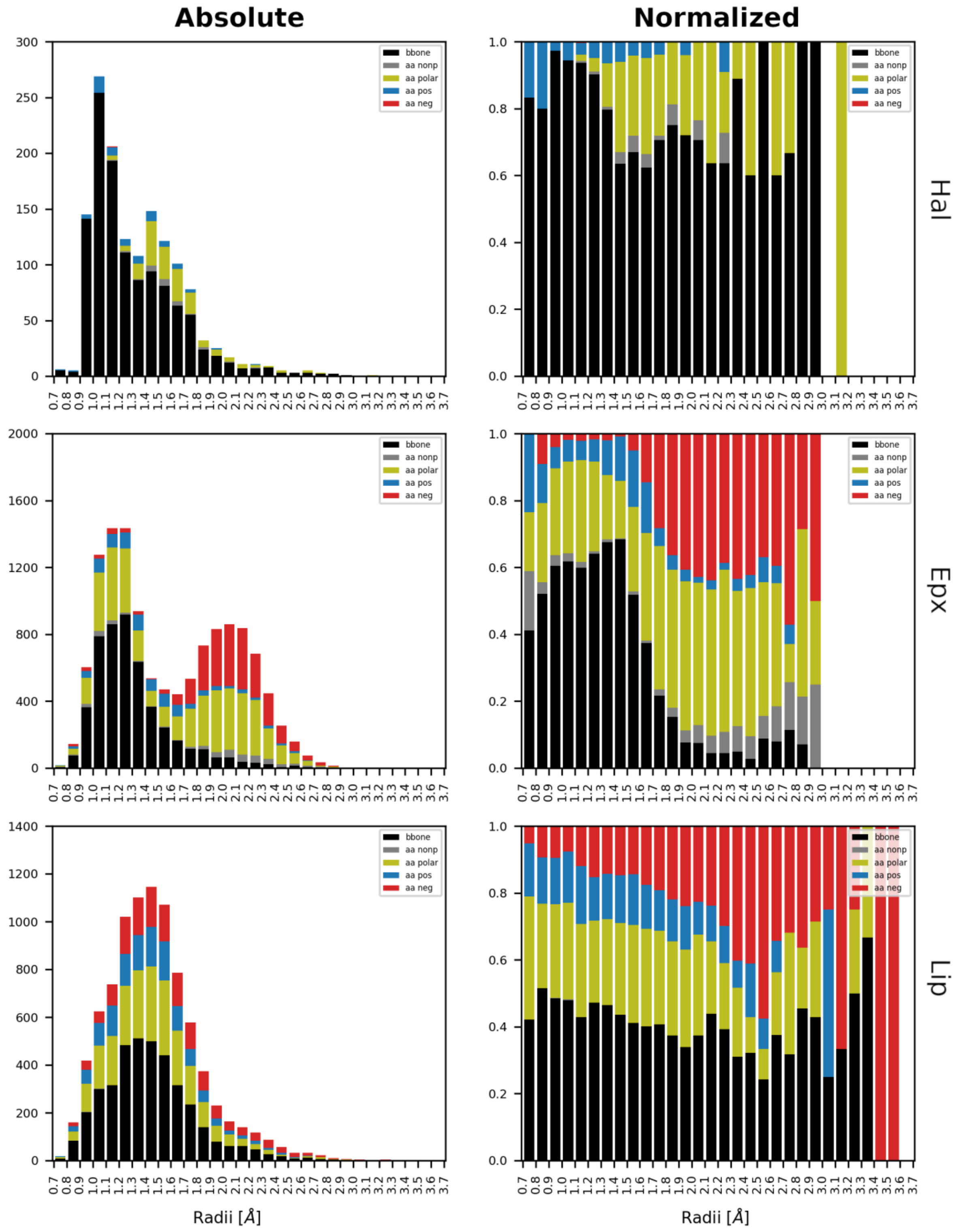


**Figure S15. Distribution of atoms involved in H-bonds at *minimal event sphere* radii in Hal, Epx, and Lip.** Histograms of the radius at which a water event occurs are presented in absolute and normalized scales for Hal, Epx, and Lip. Colors represent the type of atom involved in the H-bond present in each transport event.

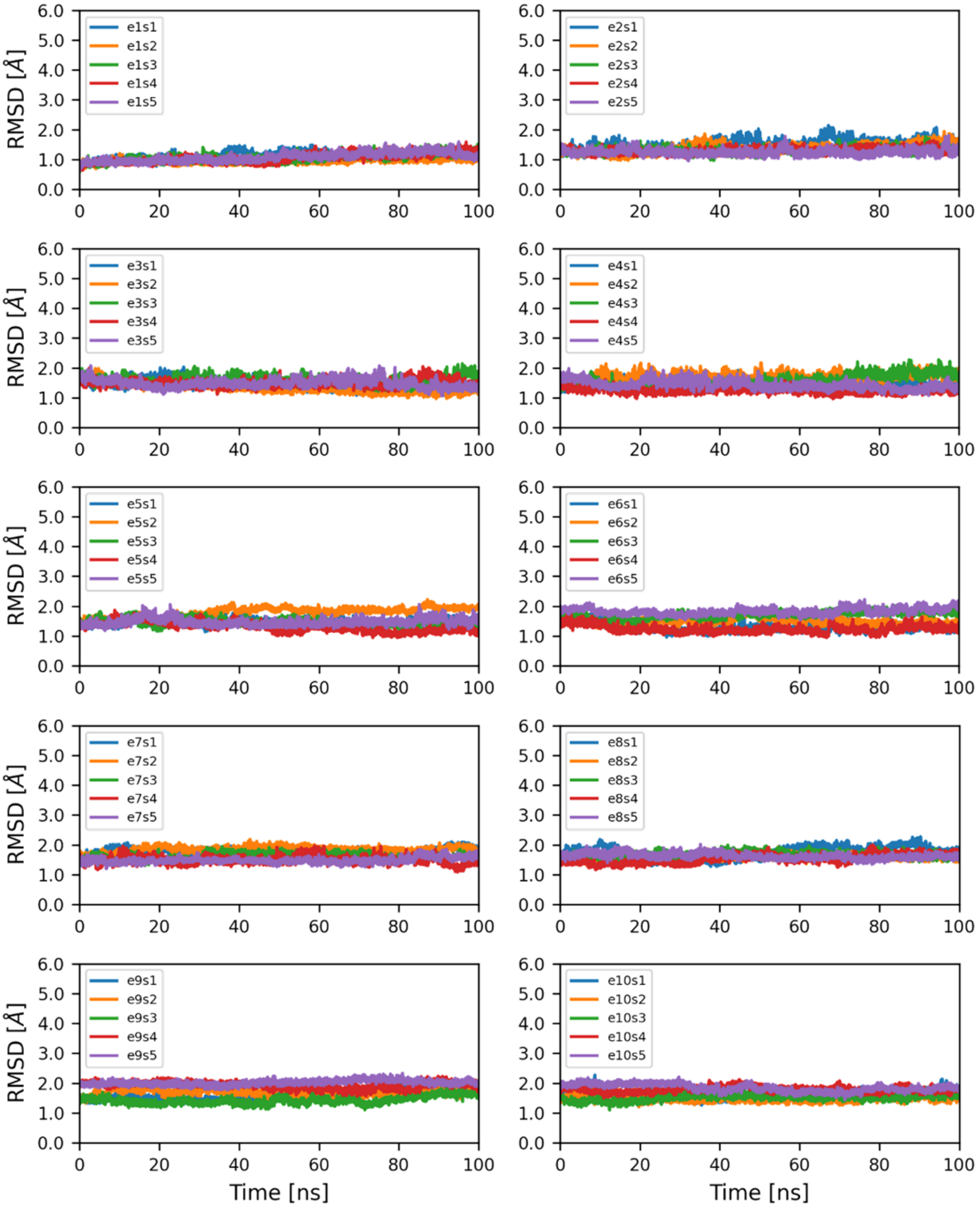


**Figure S16. RMSD of hEpx in adaptive MD simulations**. 50 MD simulations of hEpx grouped by epochs.

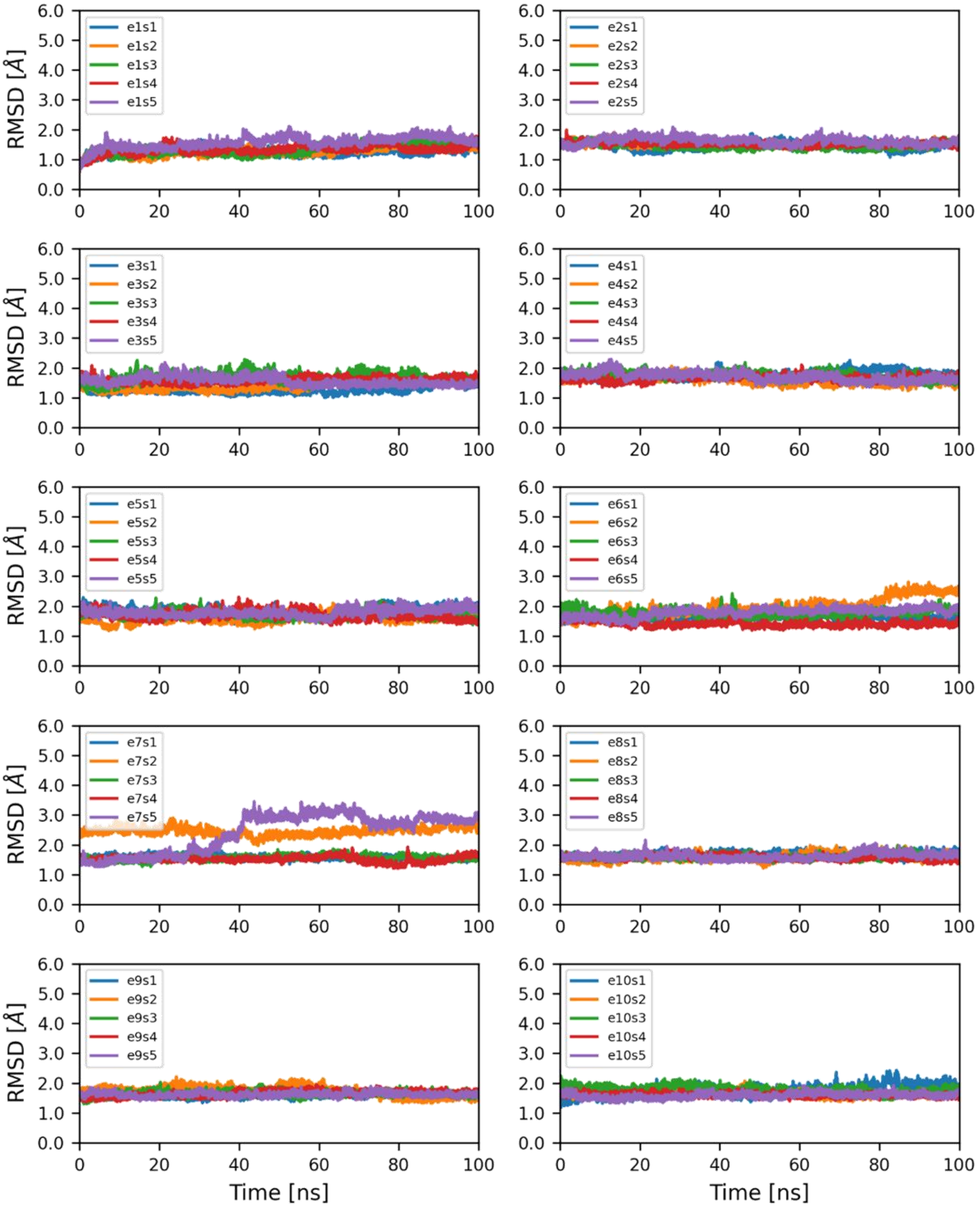


**Figure S17. RMSD of E470G in adaptive MD simulations**. 50 MD simulations of E470G grouped by epochs.

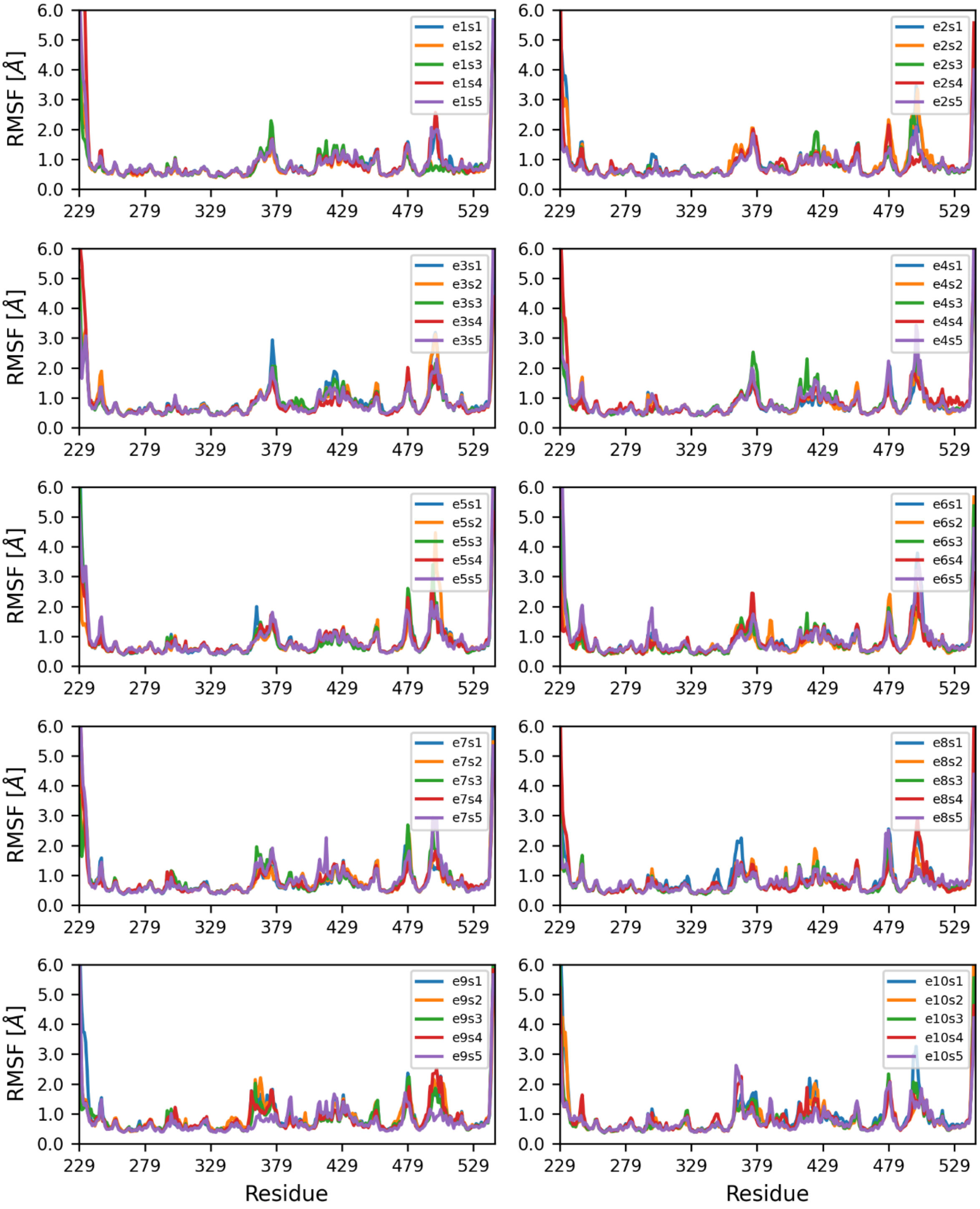


**Figure S18. RMSF of hEpx in adaptive MD simulations**. 50 MD simulations of hEpx grouped by epochs.

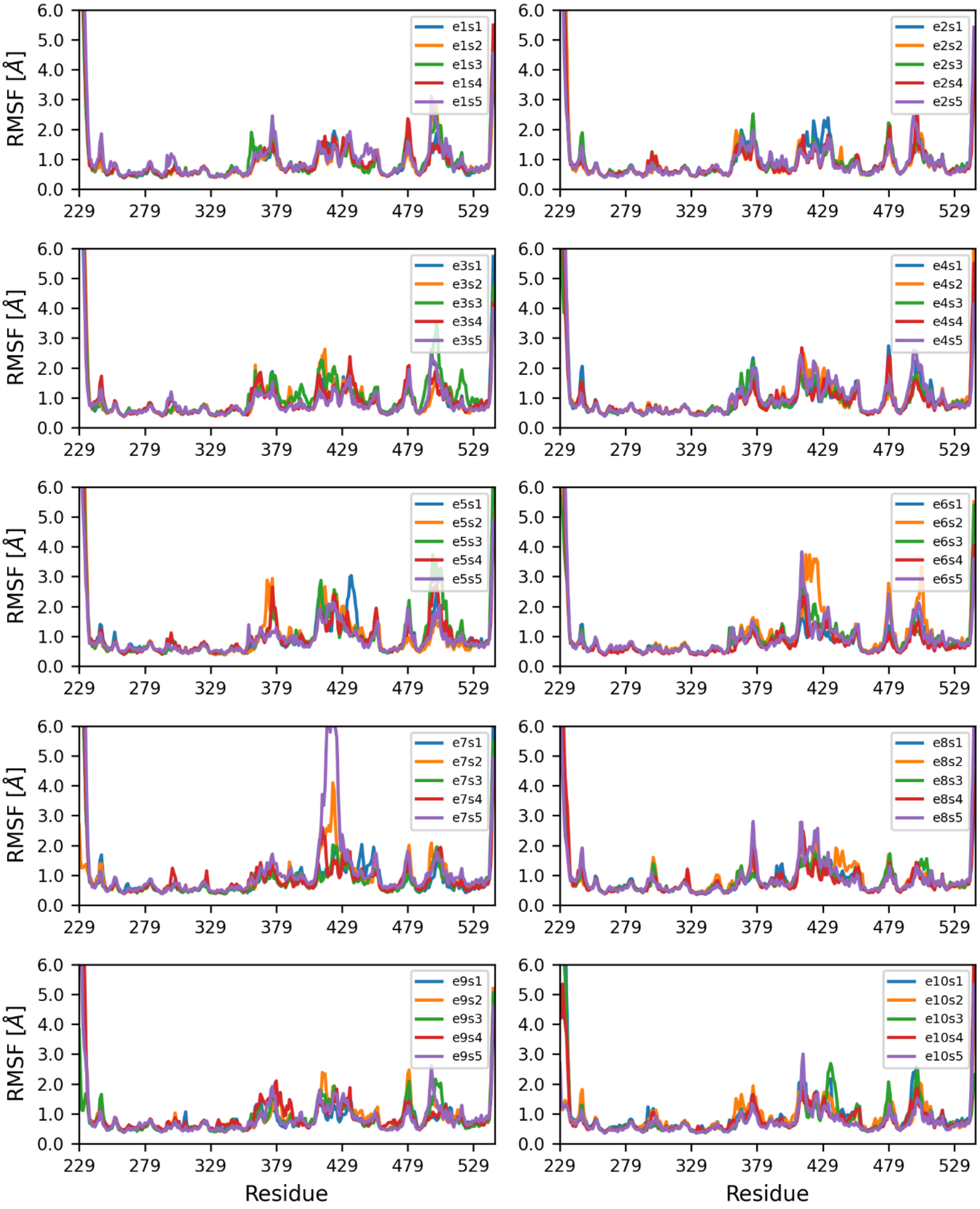


**Figure S19. RMSF of E470G in adaptive MD simulations**. 50 MD simulations of E470G grouped by epochs.

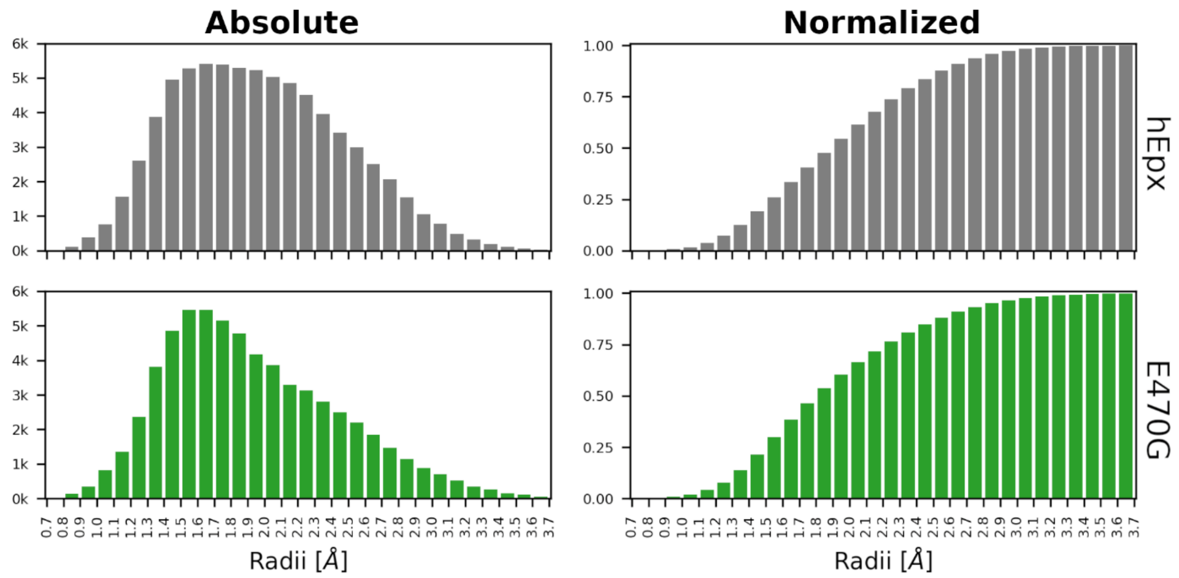


**Figure S20. Distribution of *minimal event sphere* radii for water transport in hEpx and E470G**. The absolute number of transport events and the corresponding normalized cumulative distribution, respectively.

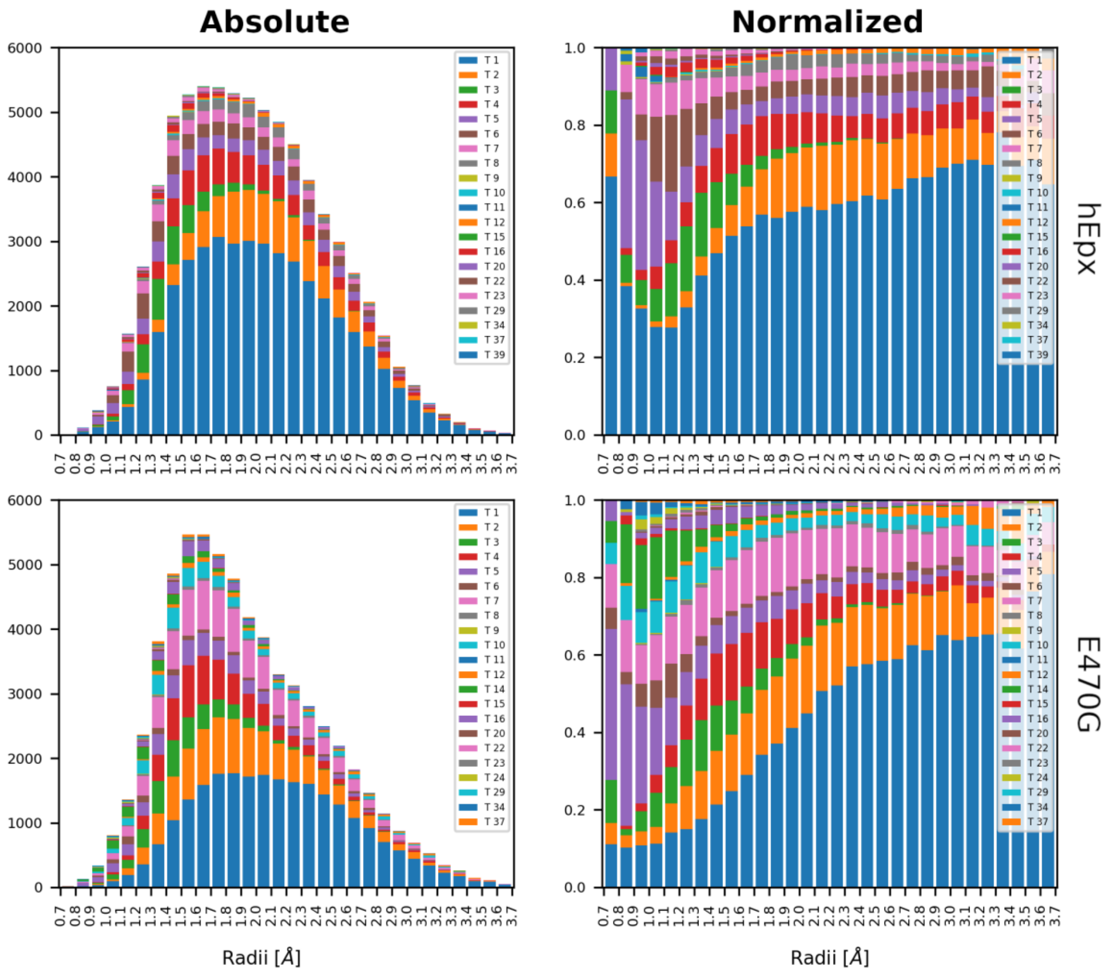


**Figure S21. Distribution of *minimal event sphere* radii in hEpx and E470G by tunnels**. Histograms of the radius at which a water event occurs are presented in absolute and normalized scales for hEpx and E470G. Colors represent individual clusters. However, it loops in due to the high number of tunnels identified.

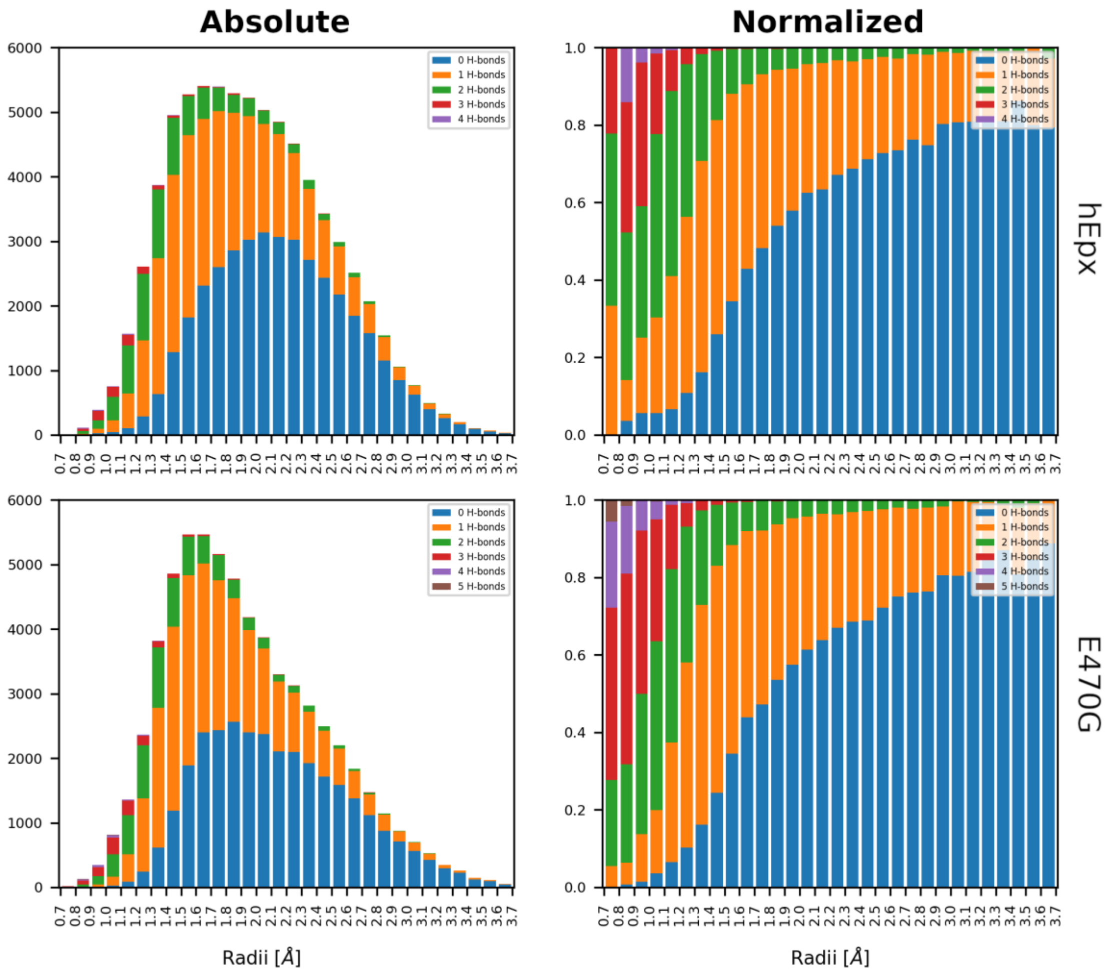


**Figure S22. Distribution of *minimal event sphere* radii in hEpx and E470G by H-bonds.** Histograms of the radius at which a water event occurs are presented in absolute and normalized scales for hEpx and E470G. Colors represent the number of H-bonds present in each transport event.

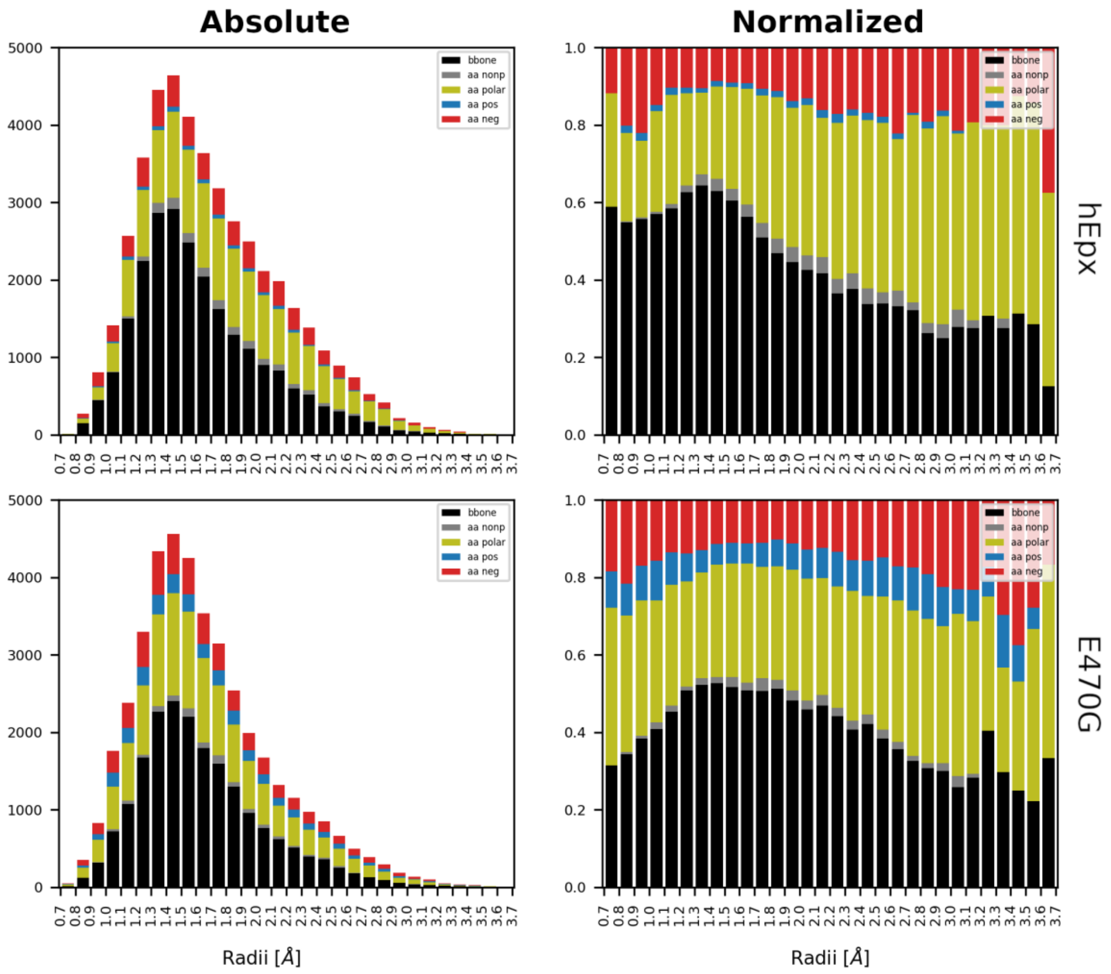


**Figure S23. Distribution of atoms involved in H-bonds at *minimal event sphere* radii in hEpx and E470G.** Histograms of the radius at which a water event occurs are presented in absolute and normalized scales for hEpx and E470G. Colors represent the type of atom involved in the H-bond present in each transport event.

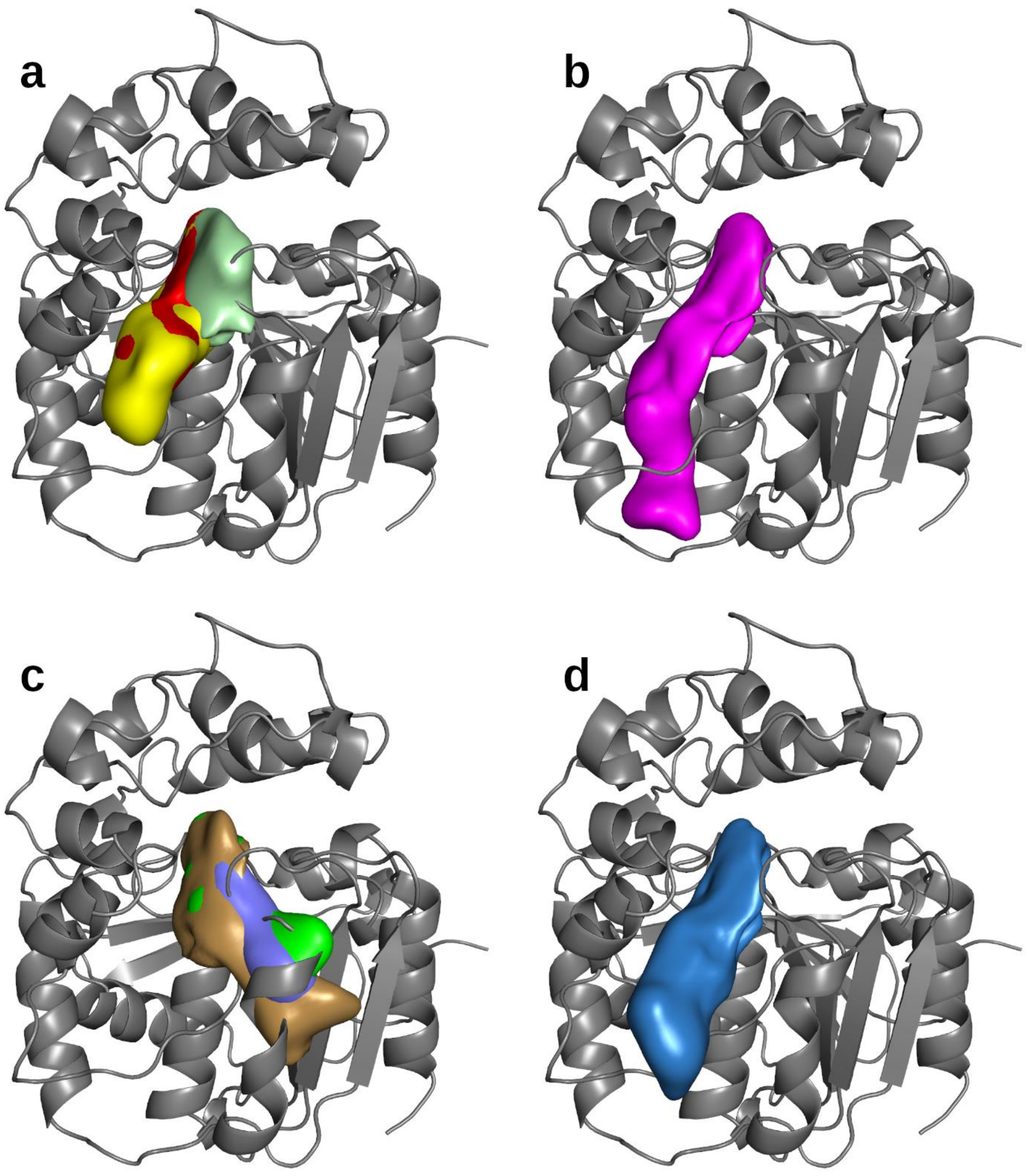


**Figure S24. Tunnels identified in hEpx and E470G.** Tunnels' densities at 20% of presence in hEpx and E470G obtained from 5 µs adaptive MD simulations. Tunnels described in the literature: a) *Tm1* (red, yellow, and pale green); b) *Tm5* (magenta); c) *Tm3* (sand, violet, and green); and d) *Tm2* (blue).

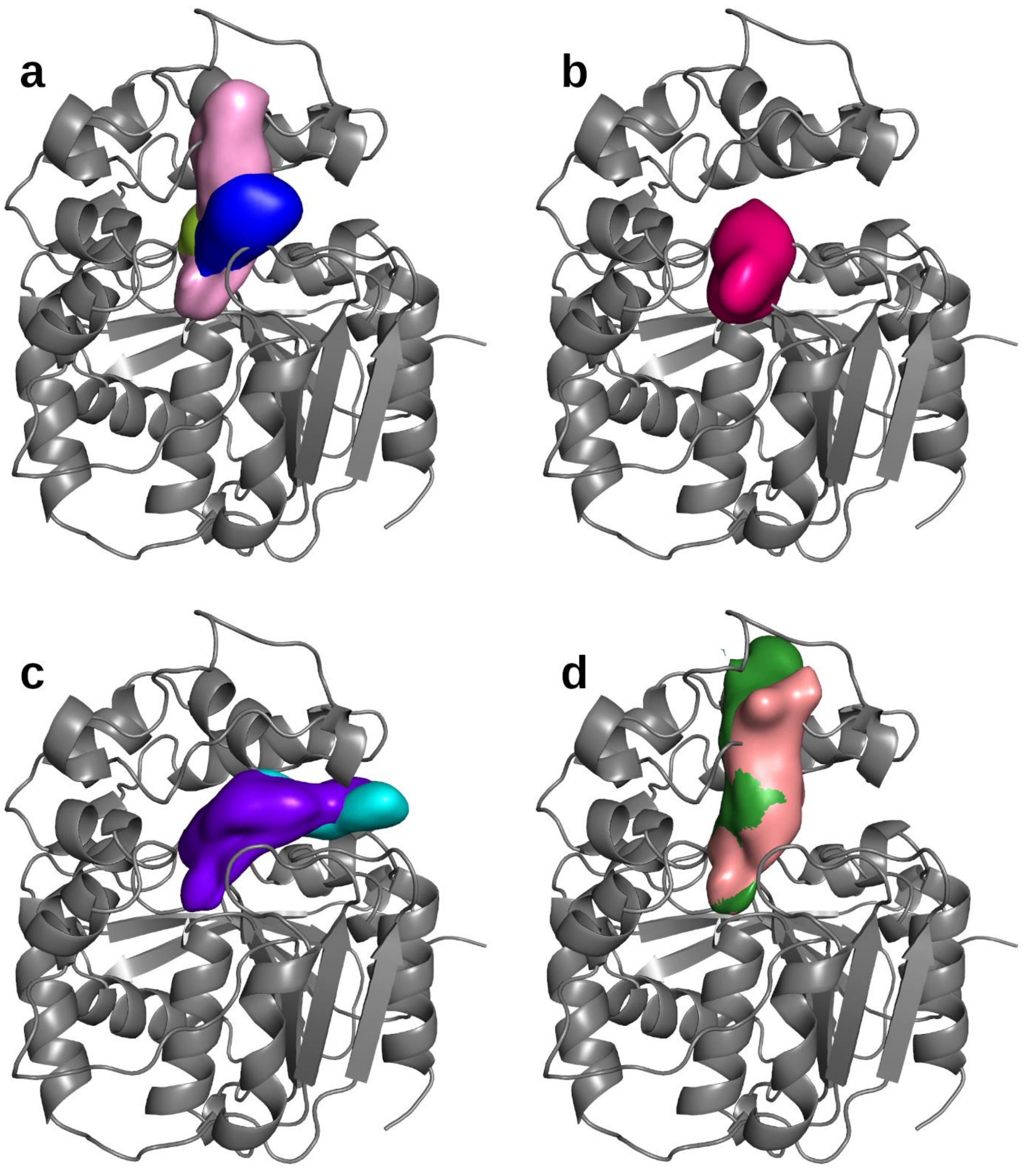


**Figure S25. Tunnels identified in hEpx and E470G.** Tunnels' densities at 20% of presence in hEpx and E470G obtained from 5 µs adaptive MD simulations. Tunnels described in the literature: a) *Tc/m* (blue, lemon, and pink); b)*Tg*; c) Tside (cyan and purple); and d) *Tcap1* (salmon and green).

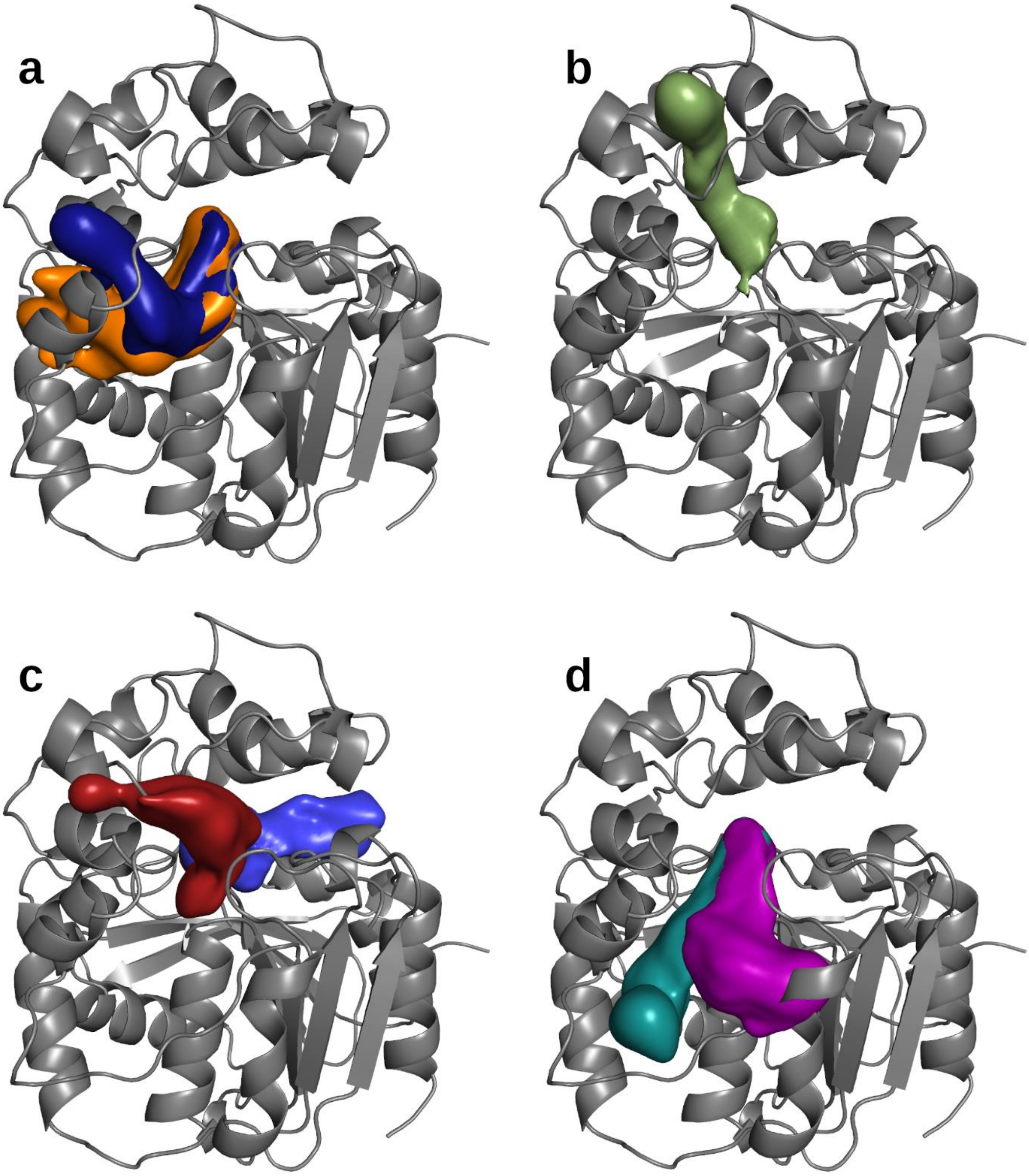


**Figure S26. Tunnels identified in hEpx and E470G.** Tunnels' densities at 20% of presence in hEpx and E470G obtained from 5 µs adaptive MD simulations. Tunnels described in the literature: a) *Tcap4* (orange and dark blue), b) *Tcap2*. Newly observed lesser tunnels that also participate in water transport c) *Tnew_A* and *Tnew_B* (in wine and blue, respectively) and d) *Tnew_C* and *Tnew_D* (in magenta and teal, respectively).

**Table S1. Tunnels identified in Hal, Epx, and Lip and their corresponding naming in the literature.**

| Protein | Tunnel | Tunnel ID | Number of water molecules transported |
|---|---|---|---|
| Hal | *p1* | 1 | 2080 |
| | *p2a* | 2 | 240 |
| | *p2b* | 30 | 6 |
| | *p2c* | 5 | 230 |
| | *p3* | 3 | 15 |
| | *new_A* | 7 | 7 |
| | *new_B* | 10 | 27 |
| Epx | *TM1* | 1, 5 | 49759 |
| | *TC/M* | 2 | 2399 |
| | *TM2* | 3 | 1907 |
| | *new_A* | 4 | 714 |
| | *new_B* | 6 | 16 |
| | *new_C* | 7 | 11 |
| | *new_D* | 8 | 6 |
| | *new_E* | 9 | 11 |
| | *new_F* | 10 | 50 |
| | *new_G* | 11 | 42 |
| | *new_H* | 12 | 39 |
| | *new_I* | 13 | 7 |
| | *new_J* | 14 | 24 |
| | *new_K* | 15 | 32 |
| | *new_L* | 16 | 6 |
| | *new_M* | 17 | 8 |
| | *new_N* | 19 | 35 |
| | *new_O* | 21 | 27 |
| | *new_P* | 22 | 145 |
| | *new_Q* | 23 | 15 |
| | *new_R* | 24 | 10 |
| | *new_S* | 25 | 25 |
| Lip | *main* | 1, 4 | 32578 |
| | *ester* | 7 | 2 |
| | *new_A* | 2 | 3448 |
| | *new_B* | 3 | 2616 |
| | *new_C* | 5 | 266 |
| | *new_D* | 6 | 89 |
| | *new_E* | 8 | 143 |
| | *new_F* | 9 | 152 |
| | *new_G* | 10 | 77 |
| | *new_H* | 11 | 2097 |
| | *new_I* | 12 | 4136 |
| | *new_J* | 14 | 417 |
| | *new_K* | 15 | 44 |
| | *new_L* | 16 | 945 |
| | *new_M* | 17 | 9 |
| | *new_N* | 18 | 62 |
| | *new_O* | 19 | 50 |
| | *new_P* | 24 | 6 |
| | *new_Q* | 26 | 207 |
| | *new_R* | 29 | 51 |
| | *new_S* | 38 | 3 |
| | *new_T* | 66 | 5 |
| | *new_U* | 70 | 14 |

**Table S2. Water events detected in Hal, Epx, and Lip and assigned to their tunnel networks.**

| Protein | Number of transport events | | |
|---|---|---|---|
| | Total | Assigned to TransportTools superclusters | Matched to a particular tunnel in the same MD simulation |
| Hal | 2658 | 2620 | 2222 |
| Epx | 55395 | 55336 | 17786 |
| Lip | 49192 | 47493 | 11086 |

**Table S3. Water usage and average bottlenecks of tunnels from hEpx and E470G.**

| Tunnel | Water usage | | | Average Bottleneck Radius | | |
|---|---|---|---|---|---|---|
| | WT | E470G | Difference | WT | E470G | Difference |
| **1** | 41543 | 25008 | **16535** | 1.904 | 1.836 | 0.068 |
| 2 | 9033 | 9482 | -449 | 1.676 | 1.689 | -0.013 |
| 3 | 3265 | 3309 | -44 | 1.223 | 1.304 | -0.081 |
| 4 | 5866 | 5684 | 182 | 1.443 | 1.460 | -0.017 |
| 5 | 4169 | 3655 | 514 | 1.089 | 1.309 | -0.220 |
| **6** | 3846 | 1060 | **2786** | 1.074 | 0.989 | 0.085 |
| **7** | 2653 | 7548 | **-4895** | 1.221 | 1.515 | -0.294 |
| **8** | 1820 | 515 | **1305** | 1.663 | 1.652 | 0.011 |
| 9 | 22 | 26 | -4 | 0.793 | 0.797 | -0.004 |
| **10** | 160 | 2687 | **-2527** | 0.954 | 1.082 | -0.128 |
| 11 | 26 | 15 | 11 | 0.818 | 0.777 | 0.041 |
| 12 | 571 | 898 | -327 | 1.270 | 1.191 | 0.079 |
| **14** | 0 | 1343 | **-1343** | 0.820 | 1.020 | -0.200 |
| 15 | 11 | 61 | -50 | 0.851 | 0.872 | -0.021 |
| 16 | 622 | 1572 | -950 | 1.097 | 1.259 | -0.162 |
| 20 | 152 | 182 | -30 | 0.923 | 0.938 | -0.015 |
| 22 | 213 | 104 | 109 | 1.093 | 0.975 | 0.118 |
| 23 | 462 | 121 | 341 | 1.233 | 1.117 | 0.116 |
| 24 | 0 | 113 | -113 | 0.742 | 0.884 | -0.142 |
| 29 | 132 | 60 | 72 | 1.539 | 1.767 | -0.228 |
| 34 | 81 | 316 | -235 | 1.153 | 1.108 | 0.045 |
| 37 | 63 | 288 | -225 | 1.408 | 1.369 | 0.039 |
| 39 | 44 | 0 | 44 | 0.842 | 0.791 | 0.051 |
| **Total** | **74754** | **64047** | **10707** | | | |
